# Mouse Aging Cell Atlas Analysis Reveals Global and Cell Type Specific Aging Signatures Revision 1

**DOI:** 10.1101/2019.12.23.887604

**Authors:** Martin Jinye Zhang, Angela Oliveira Pisco, Spyros Darmanis, James Zou

## Abstract

Aging is associated with complex molecular and cellular processes that are poorly understood. Here we leveraged the Tabula Muris Senis single-cell RNA-seq dataset to systematically characterize gene expression changes during aging across diverse cell types in the mouse. We identified aging-dependent genes in 76 tissue-cell types from 23 tissues and characterized both shared and tissue-cell-specific aging behaviors. We found that the aging-related genes shared by multiple tissue-cell types also change their expression congruently in the same direction during aging in most tissue-cell types, suggesting a coordinated global aging behavior at the organismal level. Scoring cells based on these shared aging genes allowed us to contrast the aging status of different tissues and cell types from a transcriptomic perspective. In addition, we identified genes that exhibit age-related expression changes specific to each functional category of tissue-cell types. All together, our analyses provide one of the most comprehensive and systematic characterizations of the molecular signatures of aging across diverse tissue-cell types in a mammalian system.

## Introduction

Aging leads to the functional decline of major organs across the organism and is the main risk factor for many diseases, including cancer, cardiovascular disease, and neurodegeneration^34,42^. Past studies have highlighted different hallmarks of the aging process, including genomic instability, telomere attrition, epigenetic alterations, loss of proteostasis, deregulated nutrient sensing, mitochondrial dysfunction, cellular senescence, stem cell exhaustion, and altered intercellular communication^5,34,43,63^. However, the primary root of aging remains unclear, and the underlying molecular mechanisms are yet to be fully understood.

To gain a better insight into the mammalian aging process at the organismal level, the Tabula Muris Consortium, which we are members of, created a single cell transcriptomic dataset called Tabula Muris Senis (TMS)^49^. TMS is one of the largest expert-curated single-cell RNA sequencing (scRNA-seq) datasets, containing over 300,000 annotated cells from 23 tissues and organs of male and female mice (Mus musculus). The cells were collected from mice of diverse ages, making this data a tremendous opportunity to study the genetic basis of aging across different tissues and cell types. The TMS data is organized into scRNA-seq expression of different tissue-cell type combinations (e.g., B cells in spleen) via expert annotation and clustering.

The original TMS paper explored primarily the cell-centric effects of aging, aiming to characterize changes in cell-type composition within different tissues. Here we provide a systematic gene-centric study of gene expression changes occurring during aging across different cell types. The cell-centric and gene-centric perspectives are complementary, as the gene expression can change within the same cell type during aging, even if the cell type composition in the tissue does not vary over time.

Our analysis focused on the TMS FACS data (acquired by cell sorting in microtiter well plates followed by Smart-seq2 library preparation^48^) because it has more comprehensive coverage of tissues and cell types (Supp. Table 1) and is more sensitive at quantifying gene expression levels as compared to the TMS droplet data. As shown in Fig. 1A, the FACS data was collected from 16 C57BL/6JN mice (10 males, 6 females) with ages ranging from 3 months (20-year-old human equivalent) to 24 months (70-year-old human equivalent). It contains 120 cell types from 23 tissues, totaling 110,096 cells. We also used the TMS droplet data (derived from microfluidic droplets) for those tissues for which the data is available, to further validate our findings on an additional dataset generated by a different method.

**Figure 1.**
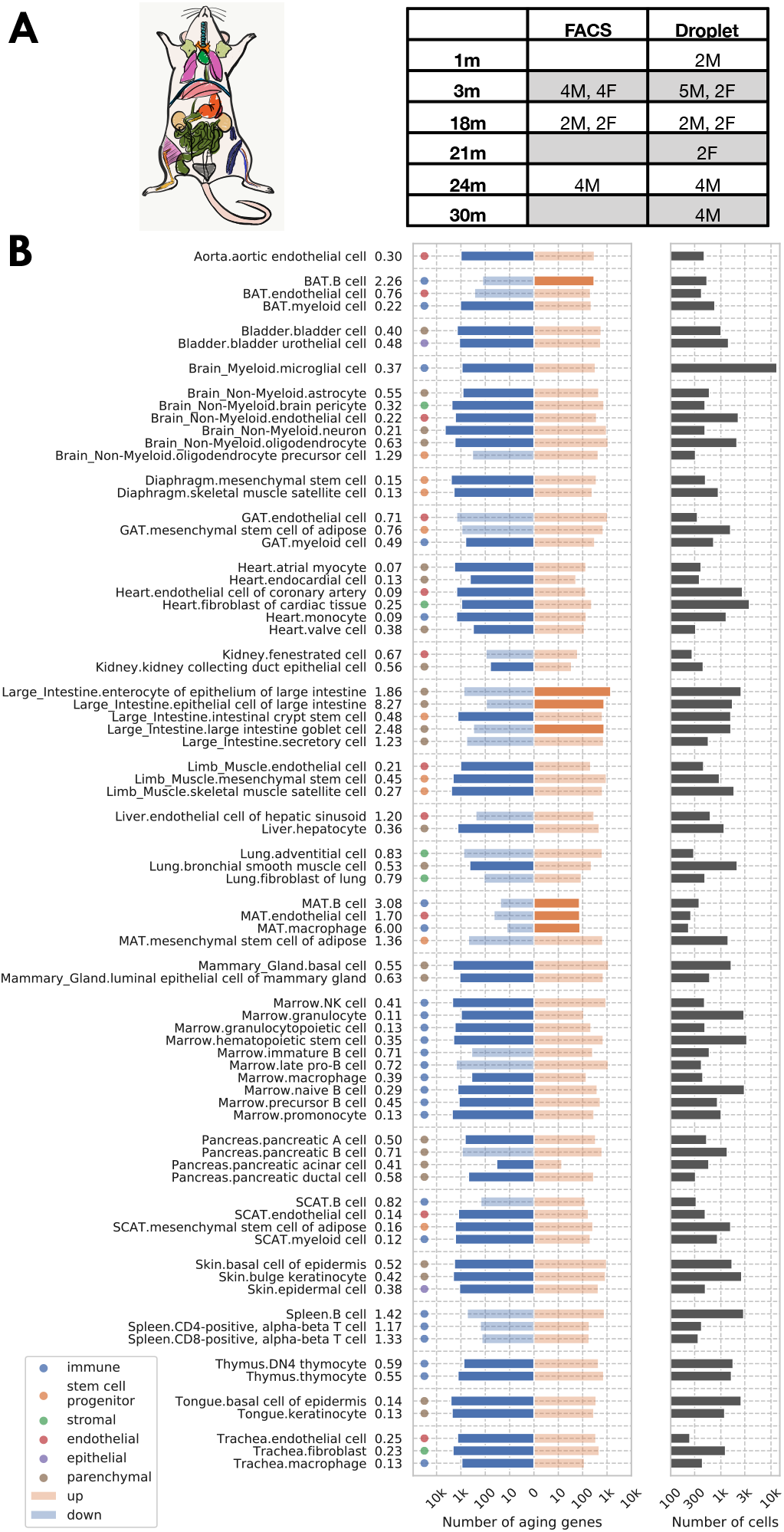
Analysis overview. A: Sample description. The TMS FACS data was collected from 16 C57BL/6JN mice (10 males, 6 females) with ages ranging from 3 months (20-year-old human equivalent) to 24 months (70-year-old human equivalent). B: Significantly aging-dependent genes in all 76 tissue-cell types in the FACS data. The left panels show the number of aging-related genes (discoveries) for each tissue-cell type, broken down into the number of up-regulated genes (orange), and the number of down-regulated genes (blue), with the numbers on the left showing the ratio (up/down). Tissue-cell types with significantly more up-/down- regulated genes (ratio>1.5) are highlighted in solid color. Most tissue-cell types have significantly more down-regulated aging genes. The right panel shows the number of cells sequenced for each tissue-cell type.

We investigated the comprehensive expression signatures of aging across tissues and cell types in the mouse. We performed systematic differential gene expression (DGE) analysis to identify aging-related genes in 76 tissue-cell type combinations across 23 tissues (Fig. 1B, Supp. Table 1). Furthermore, we characterized both shared and tissue-cell-specific aging signatures. Our study identified global aging genes, namely genes whose expression varies substantially with age in most (>50%) of the tissue-cell types. Interestingly, the expression changes of these genes are highly concordant across tissue-cell types and exhibit strong bimodality, i.e., these genes tend to be either down-regulated during aging in most of the tissue-cell types or up-regulated across the board. We leveraged this coordinated dynamic to construct an aging score based on the global aging genes. We found that the aging score is predictive of the chronological age, both in the FACS data and in multiple independent datasets. Moreover, the aging score contrasts the aging status of tissue-cell types with different functionalities and turnover rates, shedding light on the heterogeneous aging process across the 76 tissue-cell types. The score distinguished itself by its single-cell resolution and large data scale, as previous works either studied the biological age at an individual-level^12,17,19,20,22,46,47^ or focused on a small number of organs^9,44^. Overall, our analysis highlights the power of scRNA-seq in studying aging and provides a comprehensive catalog of aging-related gene signatures across diverse tissue-cell types.

## Results

### Identification of aging-related genes

We considered 76 tissue-cell types in the TMS FACS data, 26 tissue-cell types in the TMS droplet data, and 17 tissues in an accompanying bulk RNA-Seq mouse aging study^55^ (referred to as the bulk data) with sufficient sample size. We performed differential gene expression (DGE) analysis for each tissue-cell type separately, treating all cells from the tissue-cell type as samples. We test if the expression of each gene is significantly related to aging using a linear model treating age as a numerical variable while controlling for sex. We applied an FDR threshold of 0.01 (the number of comparisons corresponds to the number of genes in the tissue-cell type) and an age coefficient threshold of 0.005 (in the unit of log fold change per month, corresponding to 10% fold change from 3m to 24m). For details, please refer to the differential gene expression analysis subsection in Methods.

As shown in Fig. 1B, the number of significantly age-dependent genes per tissue-cell type ranges from hundreds to thousands. Interestingly, most tissue-cell types have more down-regulated aging-related genes than up-regulated aging-related genes, suggesting a general decrease in gene expression over aging. This down-regulation pattern is unlikely to be confounded by technical factors such as sequencing depth because the 18m/24m mice were sequenced deeper (Supp. Figs. 1C-E). By doing separate DGE analyses using mice from one sex or mice from a subset of age groups (3m/18m), we further found that such a down-regulation pattern was mostly driven by 24m mice and was not specific to one sex (Supp. Fig. 3). We also observed a similar pattern in the droplet data (Supp. Fig. 4).

In addition, we found that most aging-related genes identified in the analysis have monotonic aging trajectories, meaning that their expressions either increased or decreased monotonically during aging (Supp. Figs. 5-6). However, a subset of genes in the FACS data (13%), while up-regulated during aging overall, increased from 3m to 18m and slightly decreased from 18m to 24m; those genes are enriched in brown adipose tissue (BAT) B cells, large intestine epithelial cells, and mesenchymal adipose tissue (MAT) mesenchymal stem cells of adipose (Supp. Fig. 5C).

### Bimodal effects of aging and global aging genes

We found that most genes are significantly related to aging in at least one tissue-cell type (13,376 in the TMS FACS data and 6,233 in the TMS droplet data), consistent with the intuition that aging is a highly complex trait involving many biological processes. The aging related genes discovered in the FACS data significantly overlap with other important gene sets, including both known human and mouse aging markers as recorded in the GenAge database^60^, senescence genes^5^, transcription factors, eukaryotic initiation factors, and ribosomal protein genes (Supp. Fig. 7). Some of the top overlapping genes, significantly related to aging in most tissue-cell types, include known mouse aging markers *Jund, Apoe,* and *Gpx4* and known human aging markers *Jund, Apoe, Fos,* and *Cdc42* from the GenAge database^60^, and senescence genes *Jund, Junb, Ctnnb1, App, Mapk1* (Supp. Figs. 7A-C). In addition, we found that each tissue-cell type has around 5% aging-related genes that are shared by the GenAge human aging markers. However, we did not find any tissue-cell types that are specifically enriched with these known human aging markers, suggesting that the conservation between mouse aging and human aging is relatively uniform across tissue-cell types (Supp. Fig. 8).

We visualized all aging-related genes (significant in ≥1 tissue-cell type) in Fig. 2A, where the color indicates the number of genes. The x-axis shows the weighted proportion of tissue-cell types (out of 76 tissue-cell types) where the gene is significantly related to aging, while the y-axis shows the weighted proportion of tissue-cell types where the gene is up-regulated. The tissue-cell-type weights used here are inversely proportional to the number of cell types in the tissue, in order to ensure equal representation of the tissues. The visualization makes it clear that there are more down-regulated aging genes than up-regulated ones, consistent with Fig. 1B. Perhaps more strikingly, a bimodal pattern is apparent, in the sense that the aging-related genes tend to have a consistent direction of change during aging across most tissue-cell types. Interestingly, it was also recently reported in other studies that many shared aging-related genes exhibit consistent direction of change during aging across mouse tissues and cell types, including the brain^66^, kidney, lung, and spleen^23^.

**Figure 2.**
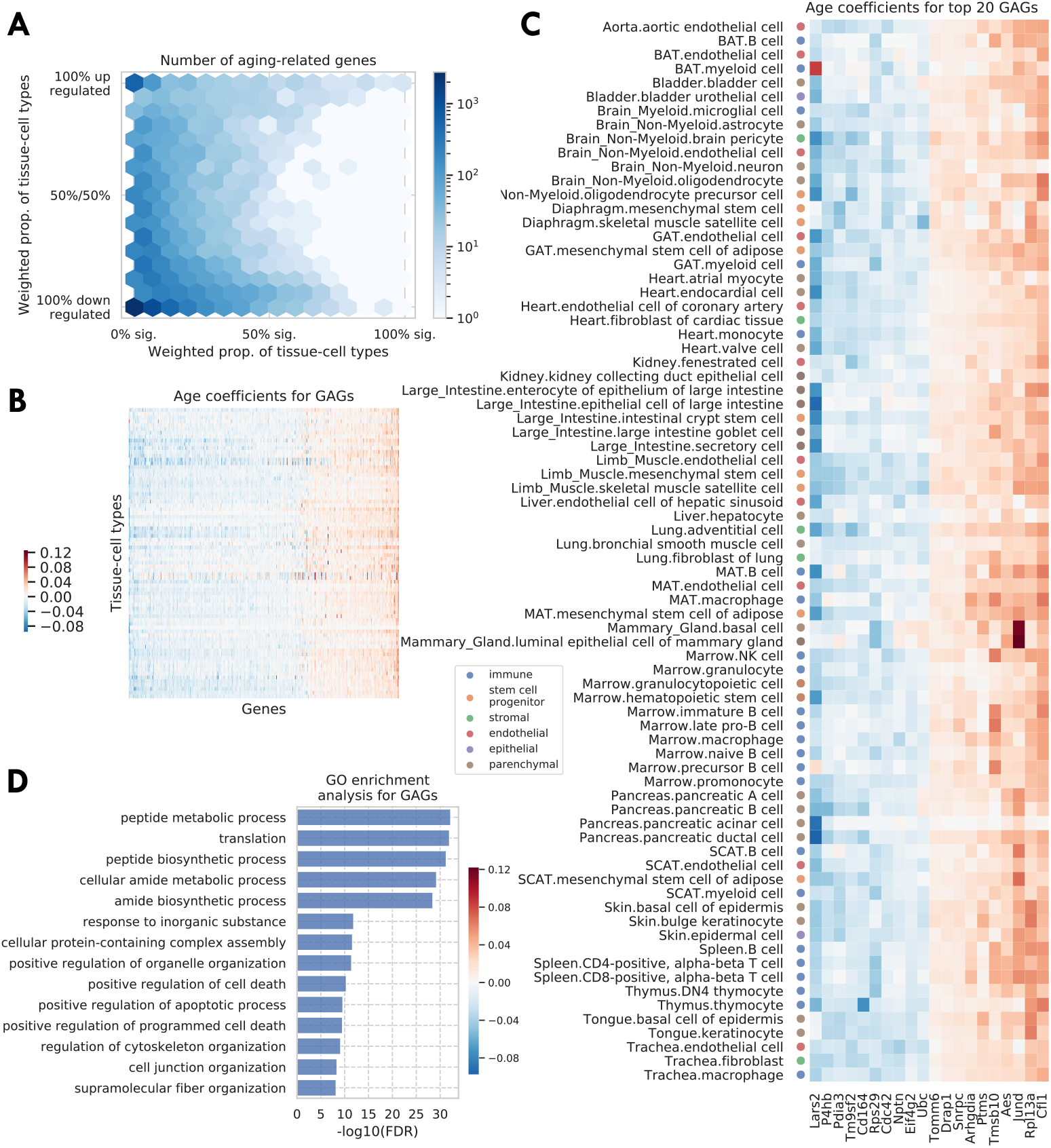
Tissue-cell level global aging genes. A: Tissue-cell level aging-related genes with color indicating the number of the genes. The x-axis shows the weighted proportion of tissue-cell types (out of all 76 tissue-cell types) where the gene is significantly related to aging, while the y-axis shows the weighted proportion of tissue-cell types where the gene is up-regulated. B-C: Heatmap of age coefficients of the global aging genes (panel B) and of the top 20 global aging genes (panel C). The age coefficients are in the unit of log fold change per month and blue/red represent down-/up- regulation. D: Top GO biological pathways for the global aging genes.

Genes that exhibit age-related expression changes across many cell types have particularly strong bimodal behavior. This motivated us to define *global aging genes* (GAGs), namely the genes that are significantly related to aging in more than 50% of weighted tissue-cell types (i.e., cell types after normalizing by cell type frequencies using the tissue-cell-type weights as described above). We identified 330 GAGs in total, among which 93 are consistently up-regulated and 190 consistently down-regulated (>80% of weighted tissue-cell types); only 47 have an inconsistent directionality (up-regulated in 20%-80% of weighted tissue-cell types). We found that GAGs significantly overlap with genes known in aging-related diseases, including strong overlap with genes related to Alzheimer’s disease (p=2.4e-11), neuroblastoma (p=1.4e-7), fibrosarcoma (p=3.3e-5), and osteoporosis (p=1.5e-4), and relatively weaker overlap with genes related to Huntington’s disease (p=2.5e-3), skin carcinoma (p=3.6e-3), kidney cancer (p=1.2e-3), acute promyelocytic leukemia (p=3.2e-3), acute myeloid leukemia (p=3e-3), endometrial cancer (p=1.6e-3), and hypertension (p=1.9e-3) (please see the global aging genes subsection in Methods for details). Our results are not sensitive to the specific choice of the 50% threshold for selecting GAGs; using different thresholds produced similar GO enrichment analysis results (Supp. Fig. 10C) or GAG scores (Supp. Figs. 13A-D) as detailed below.

We visualized the age coefficients for 10 top up-/down- regulated GAGs (a consistent direction in >80% of weighted tissue-cell types) that are related to aging in the most number of tissue-cell types, as shown in Fig. 2C. Many of these genes have been previously shown to be highly relevant to aging. For example, the down-regulation of *Lars2* has been shown to result in decreased mitochondrial activity and increase the lifespan for *C. elegans*^32^. On the other hand, *Jund* is a proto-oncogene known to protect cells against oxidative stress and its knockout may cause a shortened lifespan in mice^30^. Moreover, *Rpl13a* was observed to be up-regulated in almost all tissue-cell types. As a negative regulator of inflammatory proteins, *Rpl13a* contributes to the resolution phase of the inflammatory response, ensuring that the inflamed tissues are completely restored back to normal tissues. It also contributes to preventing cancerous growth of the injured cells caused by prolonged expression of the inflammatory genes^38,67^. Therefore, it is interesting to observe the up-regulation of *Rpl13a* given that most old mice have severe inflammatory symptoms.

As shown in Fig. 2D, Gene Ontology (GO) biological pathway enrichment analysis revealed that the 330 GAGs are associated with apoptosis, translation, biosynthesis, metabolism, and cellular organization. These biological processes are highly relevant to aging^1,2,62^ and are shared across most cell types, consistent with the intuition that GAGs represent the global aging process across tissue-cell types. In addition, the KEGG pathways associated with the GAGs are consistent with the GO terms and additionally highlighted immune-related pathways and multiple aging-related diseases (Supp. Fig. 9A). Moreover, the findings were supported by the similar analyses on the set of 59 GAGs discovered in the droplet data (Supp. Figs. 10A-B). We also performed pathway enrichment analysis using the Ingenuity Pathway Analysis software (IPA)^26^, which confirmed our findings for the biological processes associated with the GAGs (Supp. Fig. 9B). Of note is the finding that the mTOR pathway, a known aging associated pathway, is predicted to be inhibited given the expression of the GAGs^21,45,64^ (Supp. Fig. 9C). Interestingly, mTOR down-regulation has been shown to promote longevity^29,57^, a further indication that the GAGs are related to the aging process.

### GAG score contrasts the heterogeneous aging status of tissue-cell types

Following the analysis of global aging genes, we next leveraged these marker genes to characterize the holistic aging status of different tissue-cell types. We aggregated the expression of global aging markers into a single score for each cell, referred to as the GAG score (global aging gene score, please see the GAG score subsection in Methods for details). We used the FACS data to identify the GAGs because it has more comprehensive coverage of different tissue-cell types. Intuitively, the GAG score tags the global aging process, and it reflects both the chronological age of the organism as well as the tissue-cell-type specific aging effects. To formally dissect these components, we defined a fixed-effect model with the GAG score being the response variable and various other factors, including the chronological age, sex, and binary-coded tissue-cell types, being explanatory variables. As a sanity check, the chronological age effect on the GAG score is significantly positive (p<1e-100). However, while most young cells have smaller GAG scores and old cells have larger GAG scores, we also found four tissue-cell types whose GAG scores are similar between age groups and eight tissue-cell types that have a subpopulation of cells whose GAG scores are more similar to that of cells from a different age group (Supp. Fig. 11), highlighting the heterogeneity of the aging process across tissue-cell types.

We next considered the tissue-cell-type effects on the GAG score. Intuitively, a larger GAG score effect suggests that the corresponding cell type could be molecularly more sensitive to aging compared to other cells in the same animal. As shown in Fig. 3A, immune cells and stem cells have higher GAG score effects, while most parenchymal cell types have lower GAG score effects; such a contrast is also statistically significant, as shown in Fig. 3B. Indeed, immune cells and stem cells are known to undergo the most substantial changes with aging. Specifically, the aging of the immune system is commonly linked to the impaired capacity of elderly individuals to respond to new infections^40^. Also, adult stem cells are critical for tissue maintenance and regeneration, and the increased incidence of aging-related diseases has been associated with a decline in the stem cell function^10^. On the other hand, parenchymal cells like pancreatic cells, neurons, heart myocytes, and hepatocytes have lower aging scores. This could be an indication that these tissue-specialized cell types are more resilient to aging, and are able to maintain their functions despite the changes in the animal.

**Figure 3.**
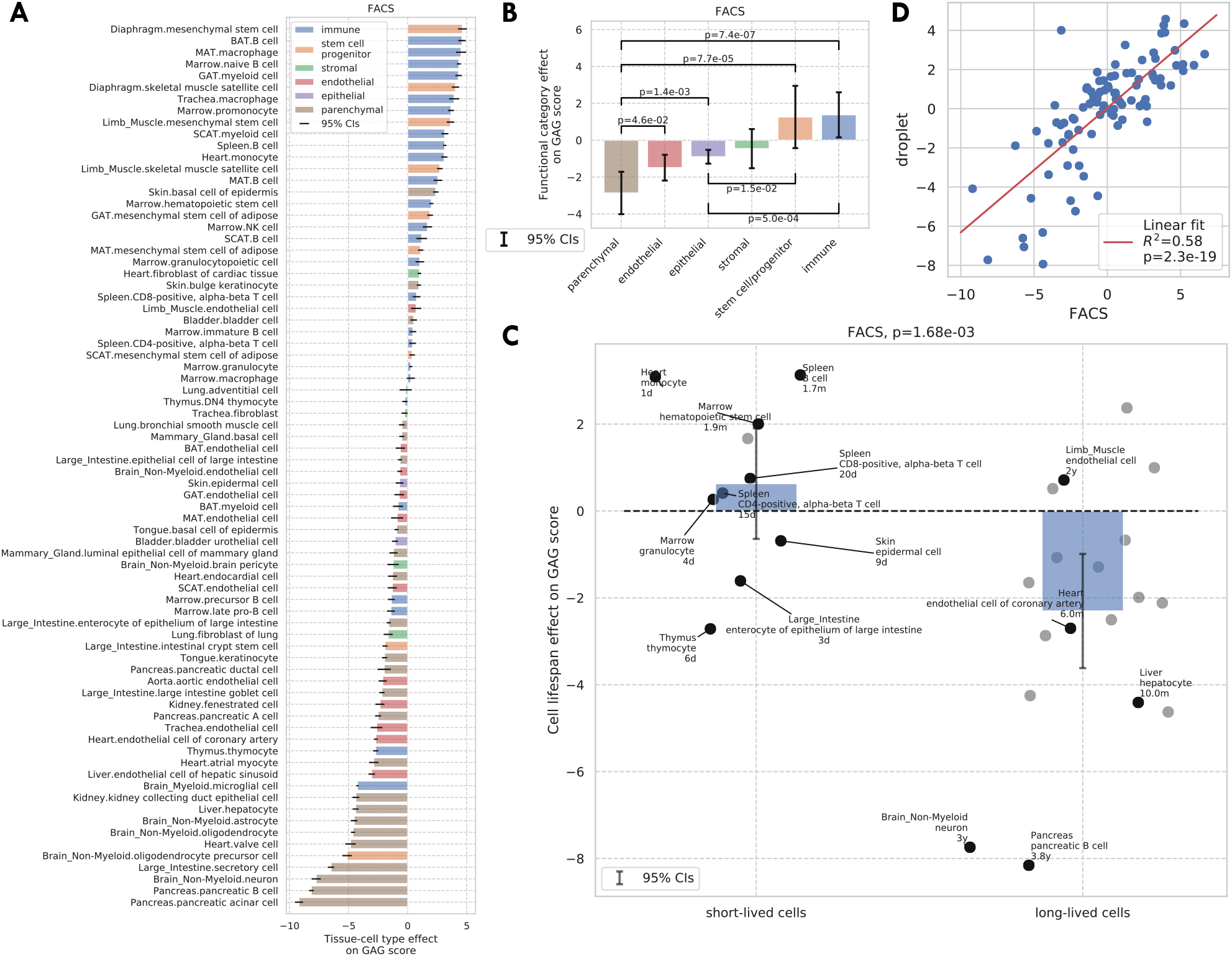
GAG score. A: Tissue-cell GAG score effects with 95% confidence intervals. The color represents the functional category of the tissue-cell type. B-C: Effects of cell functional categories (panel B) and binary cell lifespan (panel C) on the GAG score, meta-analyzed over all tissue-cell types within the group. A positive y-value means that the cells in the group (functional category group for panel B and binary cell lifespan group for panel C) have higher GAG score values than other cells of the same age and sex. 95% confidence intervals and nominal p-values are provided to quantify the differences between categories. In panel C, the average lifespan annotation is also provided for a subset of tissue-cell types where such information is available. D: Comparison between the tissue-cell GAG score effects estimated from the FACS data (x-axis) and from the droplet data (y-axis). Each dot corresponds to a tissue-cell type, and a linear fit is provided showing that the estimates are consistent.

We also found that the tissue-cell GAG score effects are in general positively correlated with the cell turnover rate. For example, short-lived cells like skin epidermal cells, monocytes, and T cells^15,24,65^ have higher GAG score effects while long-lived cells like neurons, oligodendrocytes, pancreatic *β*-cells, liver hepatocytes, and heart atrial myocytes^4,9,36,61^ have very low GAG score effects. To quantify this observation, we assigned a binary cell lifespan label to a subset of cell types where such data is available from literature^6,7,9,13,14,18,31,35,39,56,58^ (long for >180 days and short for <90 days, Supp. Table 1); using binary labels instead of the actual values allows us to incorporate more cell types whose exact lifespan information is not available but are known to be long-/short- lived. We found that short-lived cells have significantly higher GAG score effects than long-lived cells (p=1.68e-3, Fig. 3C). One possible explanation is that the GAG score is associated with the biological processes related to cell proliferation, development, and death, which are more active in cell types that have a higher turnover rate. This striking difference is also consistent with the intuition that cells that have undergone more divisions (also with higher turnover rates) are “older” and could have molecular memories.

### Validating the GAG score on external data

We performed several analyses to validate the robustness of the GAG score. First, the GAG score is not sensitive to perturbations of the current scoring method, including using different criterion to select GAGs or not performing cell-wise background correction (Supp. Figs. 13A-D). We also found that estimating tissue-cell GAG score effects using only old cells gave an almost identical result (Supp. Fig. 13H).

Next, we performed a parallel analysis to identify global aging genes on the TMS droplet data and found 59 such genes (due to smaller sample size and detection power). We found that 34 genes were shared between the droplet GAGs and the 330 FACS GAGs as described above (p=9e-48). Similar to the FACS GAGs, we also found that the droplet GAGs significantly overlap with genes known in many aging-related diseases, including Alzheimer’s disease (p=2.8e-4), neuroblastoma (p=1.2e-3), and fibrosarcoma (p=1.4e-3). In addition, we considered a set of 261 shared aging genes reported in Kimmel et al.^23^. This scRNA-seq study contains cells from the kidney, lung, and spleen in both young and old mice, and the 261 shared aging genes were defined as genes significantly related to aging in more than five cell types in the paper^23^. We found that 90 genes were shared between Kimmel et al. genes and the FACS GAGs (p=2e-105) and 42 genes were shared between Kimmel et al. genes and the droplet GAGs (p=7e-69). All p-values reported here were computed via Fisher’s exact tests.

Using the GAGs identified from the TMS FACS data, we computed the GAG score and further estimated the GAG score effects for cells in the TMS droplet data, the bulk data (treating each mouse sample as a “cell”), the Kimmel et al. data, and the dataset from Kowalczyk et al.^25^. This last dataset has only three subtypes of hematopoietic stem cells (HSCs) and was therefore omitted in other analyses. We found that the chronological age effect on the GAG score is significantly positive in all four validation datasets (p=7e-10 for the bulk data due to smaller sample size and detection power, and p<1e-100 for the other three datasets). Since the GAGs were selected based on the FACS data, the GAG score is agnostic of the age labels in the four validation datasets, confirming that the GAG score is truly indicative of the aging process.

The tissue-cell GAG score effects estimated from the other four datasets are also in line with those estimated from the FACS data (Supp. Figs. 12, 13E-F). Specifically, in the droplet data and the Kimmel et al. data, immune cell types have higher GAG score effects while epithelial, endothelial, and parenchymal cell types have lower GAG score effects. In particular for the droplet data, short-lived cell types also have higher but non-significant GAG score effects, due to a smaller number of annotated cell types. While looking at the bulk data, we found that immune-related tissues and organs such as whole blood, spleen, and marrow have the highest GAG score effects. It is interesting to observe that in the Kowalczyk et al. data, the GAG score effects of MPPs (multipotent progenitors) is less than that of ST-HSCs (short-term HSCs) which is less than that of LT-HSCs (long-term HSCs). This is exactly aligned with the differentiation potentials of these three cell types, consistent with the hypothesis that more stem-like cells have higher GAG score effects as observed in the FACS data.

Finally, we found that the tissue-cell GAG score effects are highly consistent between datasets (correlation 0.76 with p=2e-19 between the FACS data and the droplet data in Fig. 3D, correlation 0.75 with p=1e-3 between the FACS data and the Kimmel et al. data in Supp. Fig. 13G). In summary, we showed that the GAG score is capable of describing the chronological age as well as the transcriptional changes during the aging process. Furthermore, we could use the tissue-cell GAG score effects to contrast the aging status of cell types with different biological properties, including functional categories and turnover rates. This provides a comprehensive analysis demonstrating how the GAG score captures the heterogeneous molecular effects of aging.

### Category-specific aging genes

We next consider genes specific to a subset of tissue-cell types, including functional-category-specific genes, cell-type-specific genes, tissue-specific-genes, and tissue-cell-type-specific genes. Given a set of tissue-cell types, in order to have an overall meta age coefficient for cells in this tissue-cell-type set, we first combined the age coefficients of all tissue types within the set by meta-analysis; similarly, we also computed the outside-set meta age coefficient by meta-analyzing all outside-set tissue-cell types. Then we selected the genes that have significantly different within-set and outside-set meta age coefficients as the set-specific genes (please see the category-specific aging genes subsection in Methods for more details). Of note, almost no gene identified here are shared by GAGs.

In the original TMS paper^49^, each tissue-cell type was assigned one of the six functional category labels, namely the endothelial, epithelial, immune, stem/progenitor, stromal, and parenchymal cells. When examining the data by the functional category, we found that the endothelial, immune, stem, and stromal cells exhibit highly category-specific aging behavior. Indeed, we found a higher number of specific aging genes for these categories (Fig. 4B). Moreover, their age coefficients are specific to the respective functional category, as we can see from the clear block structure across tissue-cell types (Fig. 4A). In addition, when performing GO biological pathway enrichment analysis on these six sets of genes separately, we only found significant pathways for these four categories (Fig. 4C). Among them, endothelial-specific genes were associated with various processes related to angiogenesis and negative regulation of cell migration; the latter suggests decreased endothelial cell functionality during aging because endothelial cell migration is essential to angiogenesis^28^. Also, immune-specific genes were associated with activation of various immune responses, in line with a strong link between the aging process and the immune system^43,49^. In addition, stem-specific genes were associated with ossification and diverse angiogenesis processes and both stem-specific and stromal-specific genes were associated with extracellular matrix and structure organization.

**Figure 4.**
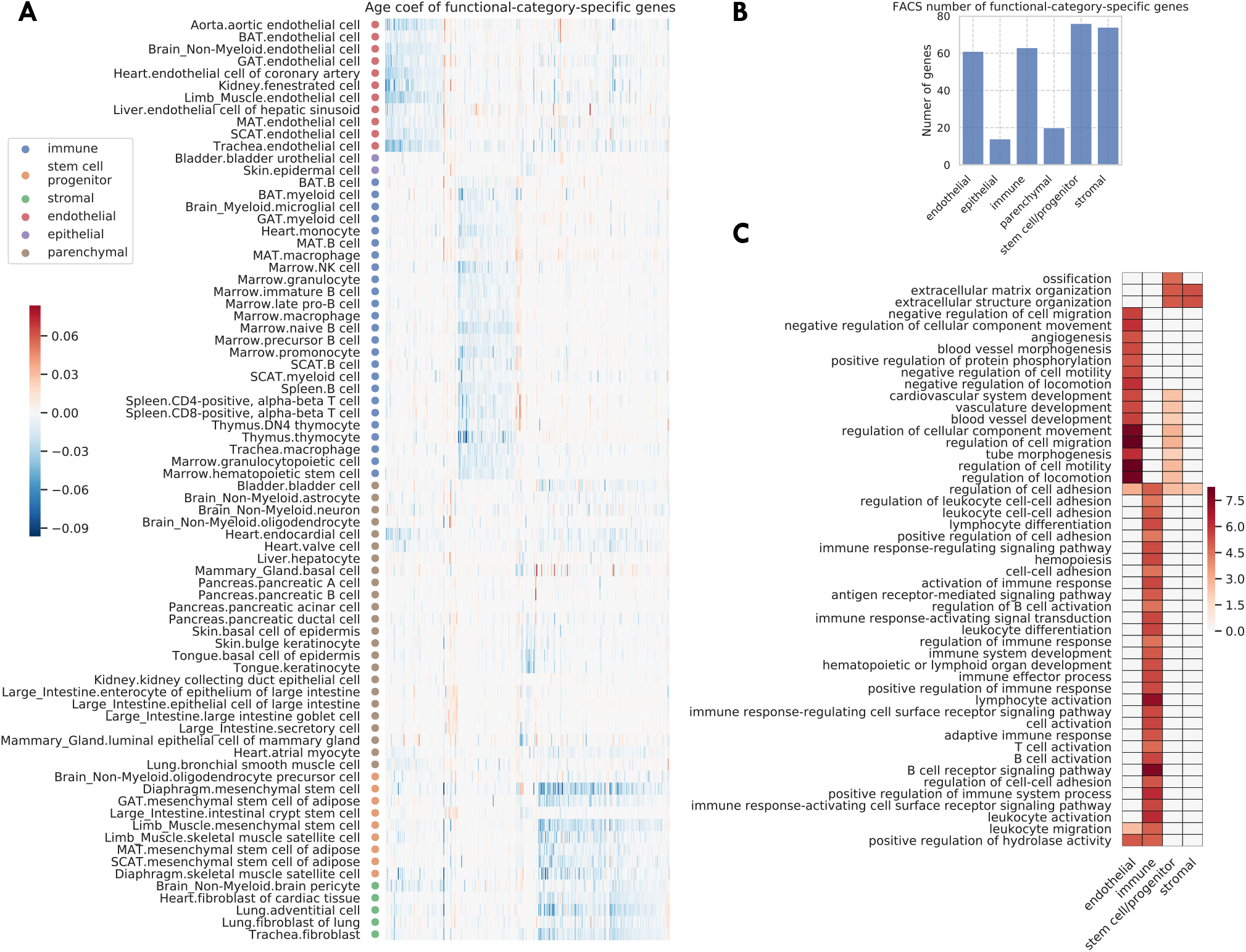
Functional-category-specific genes. A: Age coefficients, in the unit of log fold change per month, for functional-category-specific genes. Both genes in the x-axis and tissue-cell types in the y-axis are ordered by functional categories. For example, the upper-left block corresponds to endothelial-specific genes. B: Number of functional-category-specific genes for each category. C: GO biological pathways for functional-category-specific genes, with color representing the negative log10 FDR.

Such an analysis facilitates the discovery of interesting genes related to aging. For example, *C2cd4b* (Supp. Figs. 14,15A), a parenchymal-specific gene, has large age coefficients in several pancreatic, mammary gland, large intestine cell types, and almost zero age coefficients in other cell types. The increased expression of *C2cd4b* has been associated with an increased risk of type 2 diabetes^27^ and increased expression of *C2cd4b* in old mice pancreatic cell types may suggest the increased risk of type 2 diabetes for these mice. In addition, *C2cd4b* has been shown to lead to sexually dimorphic changes in body weight and glucose homeostasis^41^, in line with the fact that mammary gland is a well-known sexually dimorphic tissue. A second example is *Gsn* (Supp. Fig. 14,15B), which is down-regulated in stromal and stem cell types and up-regulated in other cell types during aging. This can be explained by its function for making gelsolin, an important protein for cell movement. Not only is cell movement important to immune and endothelial cells, but *Gsn* has also been shown to be a potential biomarker to aging-related neurodegeneration^37^.

Beyond functional-category-specific genes, we also identified genes specific to several cell types, including B cells, basal cells of epidermis, endothelial cells, macrophages, mesenchymal stem cells, mesenchymal stem cells of adipose, myeloid cells, and skeletal muscle satellite cells, corroborated by their association to related biological processes (Supp. Fig. 16). The method also allowed us to identify genes specific to each tissue. However, we did not find any genes specific to a single tissue-cell type. All the gene sets are available in Supp. Table 3.

### Tissue level analysis and validation

We repeated the aging analyses at the tissue level in order to compare and assess the robustness of our tissue-cell level findings. We carried out a DGE analysis for every tissue by pooling the cells from all cell types in the tissue. The number of discoveries for each tissue is shown in Fig. 5A. 18 out of 23 tissues have significantly more down-regulated aging genes while there are significantly more up-regulated aging genes in the large intestine, consistent with the tissue-cell level result in Fig. 1B. We also observed a similar down-regulation pattern in the droplet data in the tissue-level analysis, but a less clear pattern when looking at the bulk data (Supp. Fig. 17).

**Figure 5.**
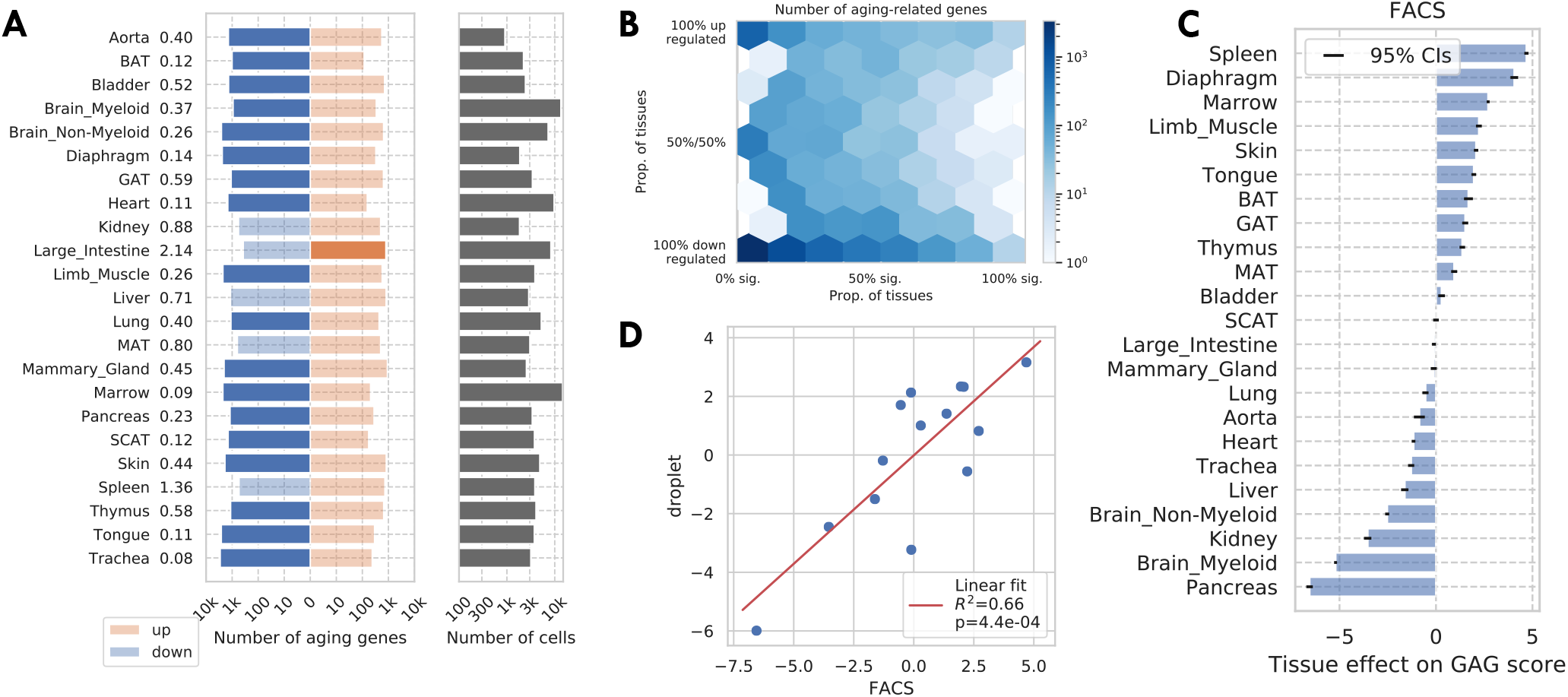
Tissue-level analysis as validation. A: Number of discoveries for each tissue. The left panels show the number of aging genes (discoveries) for each tissue, broken down into the number of up-regulated genes (orange), and the number of down-regulated genes (blue), with the numbers on the left showing the ratio (up/down). Tissue types with significantly more up-/down- regulated genes (ratio>1.5) are highlighted in the solid color. We can see most tissues have significantly more down-regulated aging genes. The right panels show the number of cells for each tissue. B: Tissue-level aging-related genes with color indicating the number of genes. The x-axis shows the weighted proportion of tissue types (out of 23 tissues) where the gene is significantly related to aging, while the y-axis shows the weighted proportion of tissues where the gene is up-regulated. C: TMS FACS tissue-level GAG score effects with 95% confidence intervals. D: Comparison between the FACS tissue-level GAG score effects and the droplet tissue-level GAG score effects. Each dot corresponds to a tissue type, and a linear fit is provided showing the estimates are consistent.

The tissue-level DGE result is summarized in Fig. 5B. Similar to the tissue-cell level analysis, we again observed a bimodal pattern, and a consistent direction of change during aging for genes shared by more tissues. We also observed a similar bimodal pattern in the bulk data (Supp. Fig. 18C). We used a threshold of 0.8 for selecting tissue-level global aging genes and found 147 such genes, out of which 138 were also tissue-cell level global aging genes, indicating a strong consistency (p<1e-100 via a Fisher’s exact test).

We next computed the tissue-level GAG score effect for each tissue using the tissue-level global aging genes. As shown in Fig. 5C, the tissue-level GAG score effects are qualitatively similar to the tissue-cell-level GAG score effects, with the immune tissues (spleen, marrow, thymus) being at the top, and brain myeloid and pancreas being at the bottom. We observed a similar consistency in the droplet data and the bulk data (Supp. Fig. 19). In addition, we observed a strong consistency between the tissue-level GAG score effects estimated from different datasets (Fig. 5D for comparison with the droplet data, Supp. Fig. 18D for comparison with the bulk data), demonstrating the robustness of the GAG score analysis.

## Discussion

This study provides a systematic and comprehensive analysis of aging-related transcriptomic signatures by analyzing 76 tissue-cell types in the TMS FACS data. Together with the analysis in the first publication of Tabula Muris Senis^49^, it forms one of the largest analysis to date of the mammalian aging process at the single-cell resolution. Of particular interest are the 330 global aging genes identified in the study. These genes exhibit aging-dependent expressions in a majority of tissue-cell types in the mouse. The global aging genes are enriched with many interesting genes, including known human and mouse aging markers, aging-related disease genes, senescence genes, transcription factors, eukaryotic initiation factors, and ribosomal protein genes. Interestingly, most of the global aging genes are strongly bimodal as their expressions either decrease or increase during aging across almost all tissue-cell types, suggesting that these genes have a uniform response to aging, which is robust to the specific tissue or cellular context. Moreover, we find a systematic decrease in expression for most genes as well as a decrease in the number of actively expressed genes, suggesting a turning off of transcription activity as the animal ages. A recent study has observed that the number of expressed genes decreases during cellular differentiation in mouse^16^. It is interesting that we quantify a similar phenomenon for aging, despite the substantial longer time-scale of aging compared to differentiation.

While we have validated our findings using the TMS droplet data, the bulk RNA-seq data, and external datasets, it is important to have further validations in future studies. In particular, the bimodal expression pattern is less apparent in the TMS droplet data, perhaps due to its limited tissue-cell-type coverage and relatively shallower sequencing depth. The remarkable bimodal consistency of the global aging genes makes them useful as biomarkers to characterize the aging status of individual cells. We proposed a new aging score, namely the GAG score, based on the global aging genes. The tissue-cell-type-specific GAG score effects quantify how sensitive each tissue-cell type is to aging and are positively correlated with the cell division rate. For example, immune cells tend to have higher GAG scores than other cells of the same age and sex, which reflects the phenomenon that they undergo many cycles of cell division and also change substantially during the animal’s lifespan. One hypothesis is that the GAG score captures some aspects of the true biological age of the cells, which could be different from the birth age of the animal. An interesting direction of future work is to further investigate this model with functional experiments. In line with this, it would be important to study how some of the transcriptomic changes we quantify here, e.g., the down-regulation of mTOR, point towards healthy aging or how can they inform experiments that can uncover the mechanism to ameliorate the aging effects.

The GAG score is also related to the transcriptome age predictors developed in previous works^12,17,19,46^, in the sense that they all use the gene expression information and are predictive of the animals’ chronological age. The commonly-used approach in previous works is to train a model (e.g., linear/logistic regression model) to predict the individuals’ chronological age from their gene expression. Instead of a model-fitting algorithm, our GAG score uses the global aging genes that were selected from a broad range of tissue-cell types in an unbiased manner, by meta-analyzing the differential gene expression (DGE) results of 76 tissue-cell types and putting each tissue-cell type on the same footing. This ensures that the genes used by the GAG score capture shared aging process and are not biased towards certain tissue-cell types. Indeed, the global aging genes were shown to be associated with biological processes that are highly relevant to aging (Fig. 2), providing better interpretability of the score. In comparison, previous studies focused on only one specific tissue, such as the blood^17,19,46^ or dermal fibroblasts^12^, and hence may have selected genes that were biased towards that particular tissue.

Overall, our study provides a comprehensive characterization of aging genes across a wide range of tissue-cell types in mice. In addition to the biological insights, it also serves as a comprehensive reference for researchers working on related topics.

## Methods

### Data preprocessing

We considered five datasets, namely the TMS FACS data, the TMS droplet data, the data in Schaum et al.^55^ (referred to as the bulk data), the data in Kimmel et al.^23^, and the data in Kowalczyk et al.^25^. For the TMS FACS data and the TMS droplet data, we filtered out genes expressed in fewer than 3 cells, filtered out cells expressing fewer than 250 genes, and discarded cells with a total number of counts fewer than 5,000 for the FACS data and a total number of unique molecular identifiers (UMIs) fewer than 2,500 for the droplet data. For the bulk data, we filtered out genes expressed in fewer than 5 samples, and filtered out samples expressing fewer than 500 genes. We did not filter cells for the other two datasets. For all five datasets, we normalized each sample to have 10,000 reads/UMIs per sample, followed by a log transformation (log(x+1) where x is the read count). We note that such a procedure is the same as that in the original paper^49^. We did not correct for batch effects because the data were centrally collected and no substantial batch effects were identified in the original TMS paper^49^.

### Differential gene expression analysis

As shown in Supp. Table 1, we considered 76 tissue-cell types in 23 tissues with more than 100 cells in both young (3m) and old (18m, 24m) age groups for the TMS FACS data; 26 tissue-cell types in 11 tissues with more than 500 cells in both young (1m, 3m) and old (18m, 21m, 24m, 30m) age groups for the TMS droplet data; and all 17 tissues for the bulk data^55^. We required more cells for the TMS droplet data than the TMS FACS data because the droplet data has a much lower sequencing depth (6000 UMIs per cell, as compared to 0.85 million reads per cell for the FACS data). Also, we did not focus on the TMS droplet data for the main results due to its limited tissue and cell type coverage.

We performed a DGE analysis for cells in each tissue-cell type separately. In the DGE analysis for a tissue-cell type, all cells in the tissue-cell type were treated as samples, and a separate test was performed for each gene with the observations being the expressions of the gene across the cells. We identified genes significantly related to aging using a linear model treating age as a continuous variable while controlling for sex, namely,

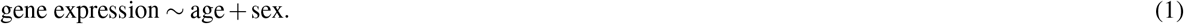

Since the tests were performed at a cell level for each tissue-cell type, the cell numbers will only affect the detection power but will not bias the result. Therefore, it is not a confounding factor and was not controlled for. We used the MAST package^11^ (version 1.12.0) in R to perform the DGE analysis. The zero counts in the scRNA-seq data were handled by the MAST package and we do not observe other types of missing data. We did not control for the cellular detection rate^11^ (CDR, corresponding to the number of expressed genes per cell) as suggested by the authors of the package because we found that CDR is positively correlated with age and negatively correlated with technical covariates such as sequencing depth and the number of detected ERCC spike-ins, both when considering all cells or considering each sex separately (Supp. Fig. 1). As a result, controlling for CDR may remove genuine aging effects. Nonetheless, we found that the age coefficients, estimated with and without CDR correction, were highly correlated (0.89 for the FACS data and 0.93 for the droplet data, Supp. Fig. 2), ruling out the possibility that CDR correction would significantly alter the result.

We used the Benjamini-Hochberg (FDR) procedure^3^ to control for multiple comparisons, where the number of comparisons corresponds to the number of genes in the tissue-cell type. We applied an FDR threshold of 0.01 and an age coefficient threshold of 0.005 (in the unit of log fold change per month, corresponding to around 10% fold change from 3m to 24m) for detecting genes significantly related to aging.

### Global aging genes

We selected a gene as a global aging age (GAG) if it is significantly related to aging in more than 50% of weighted tissue-cell types. Here, the tissue-cell-type weights are inversely proportional to the number of cell types in the tissue, in order to ensure equal representation of tissues.

For the overlap between GAGs and the genes known in aging-related diseases, we considered the top 25 aging-related diseases and obtained their related genes from the Human Disease Database (MalaCards)^50–52^. We then converted the human genes to the corresponding mouse orthologs using g:Profiler^53^ (version 1.2.2). The p-values quantifying the significance of the overlap were computed using Fisher’s exact tests.

### Pathway enrichment analysis

We used g:Profiler^53^ to perform gene ontology biological pathway enrichment analysis. We considered biological pathways with FDR smaller than 0.01. We used Gene Set Enrichment Analysis (GSEA MGSig Database)^33,59^ to perform the KEGG pathway analysis. We filtered for mouse genes and considered biological pathways with FDR smaller than 0.05. We also used the Ingenuity Pathway Analysis software (IPA)^26^ to perform canonical pathway analysis (Supp. Fig. 9). For Supp. Fig. 9A, we used an FDR threshold of 1e-5 and a z-score threshold of 0.5.

### GAG score

Given a set of global aging genes (e.g., the FACS GAGs), the cell-wise GAG score for cell *i* is computed as

1. Compute the raw GAG score as the average expression of the up-regulated GAGs (up-regulated in >80% of weighted tissue-cell types) minus the average expression of the down-regulated GAGs (down-regulated in >80% of weighted tissue-cell types), i.e.,

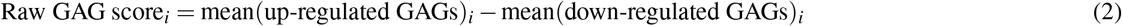
2. Following the recipe of DeTomaso et al.^8^, z-normalize the raw GAG score using the expected mean and variance of a random set of genes with the same number of up-/down- genes:

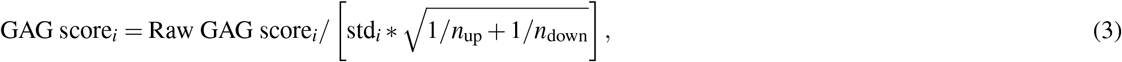

where std_*i*_ is the standard deviation of the gene expression of cell *i, n*_up_ is the number of up-regulated GAGs, and *n*_down_ is the number of down-regulated GAGs.

For estimating the GAG score effects, we model the GAG score of a cell *i* as being linearly dependant of the chronological age, sex, and tissue-cell type of the cell, namely,

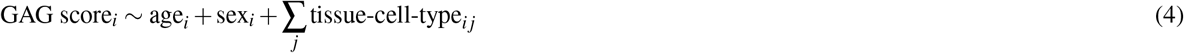

Here, age_*i*_ is the age of the animal (in months) that cell *i* comes from, sex_*i*_ is one if cell *i* comes from a male and zero otherwise, and tissue-cell-type_*ij*_ is one if cell *i* belongs to tissue-cell type *j* and zero otherwise. We do not include the intercept term because all binary coded tissue-cell types sum up to one. Finally, we further center both response and explanatory variables and perform an ordinary least square regression to estimate the GAG score effects for the age, sex, and each tissue-cell type.

Meta-analyses in Figs. 3B-C and Supp. Figs. 13E-F were performed assuming a random effect model^54^. For comparisons of the tissue-cell GAG score effects between datasets, namely those in Fig. 3D and Supp. Fig. 13G, we used all tissue-cell types instead of restricting to the 76 TMS FACS tissue-cell types and 26 TMS droplet tissue-cell types. This increased the number of overlapping tissue-cell types between datasets.

### Category-specific aging genes

We considered identifying functional-category-specific genes, cell-type-specific genes, tissue-specific genes, and tissue-cell-type-specific genes. For a set of tissue-cell types, e.g., the set of all immune tissue-cell types, the genes specific to the set (or set-specific genes) are selected as follows. For each gene, we first estimate its within-set meta age coefficient by meta-analyzing the age coefficients of all tissue-cell types in the set assuming a random effect model^54^. Specifically, it is done by assuming that there is a meta age coefficient for the set of tissue-types, and the age coefficient for each tissue-cell type in the set is a random variable whose mean is equal to the meta age coefficient of the set. Similarly, we estimate the outside-set meta age coefficient by meta-analyzing all tissue-cell types outside the set. Then, we define the set-specific genes to be genes whose

1. within-set meta age coefficient is significantly different from its outside-set meta age coefficient (FDR<0.01);
2. within-set meta age coefficient is large enough (absolute value > 0.005);
3. outside-set meta age coefficient is not significantly different from 0 (FDR>0.01).

The p-values are computed based on the mean and the standard error assuming a normal distribution, and FDR is computed with respect to all genes.

## Data availability

All data can be downloaded at https://figshare.com/articles/dataset/tms_gene_data_rv1/12827615

## Code availability

The code for reproducing all results is at https://github.com/czbiohub/tabula-muris-senis/tree/master/2_aging_signature

## Supplementary Information

1. Supplementary Figures
2. Supplementary Table 1: Summary of tissues and cell types for TMS FACS data, TMS droplet data, and the bulk data.
3. Supplementary Table 2: Significantly aging-related genes in each tissue-cell type for TMS FACS data and TMS droplet data.
4. Supplementary Table 3: Gene sets identified in the study, including GAGs and category-specific genes, for TMS FACS data and TMS droplet data.

## Acknowledgements

We would like to thank S. Quake, R. Sinha, R. Sit, J. Cool, B. van de Geijn, H. Shi, X. Xu for feedback. M.J.Z. and J.Z. are supported by NSF CCF 1763191, NIH R21 MD012867-01, NIH P30AG059307, and grants from the Silicon Valley Foundation and the Chan-Zuckerberg Initiative.

## Author contributions

M.J.Z., A.O.P., and S.D. analyzed the data. M.J.Z., A.O.P., and J.Z. wrote the manuscript. J.Z. supervised the research. All authors reviewed the manuscript.

## Supplementary Information

### Supplementary Figures

**Supplementary Figure 1.**
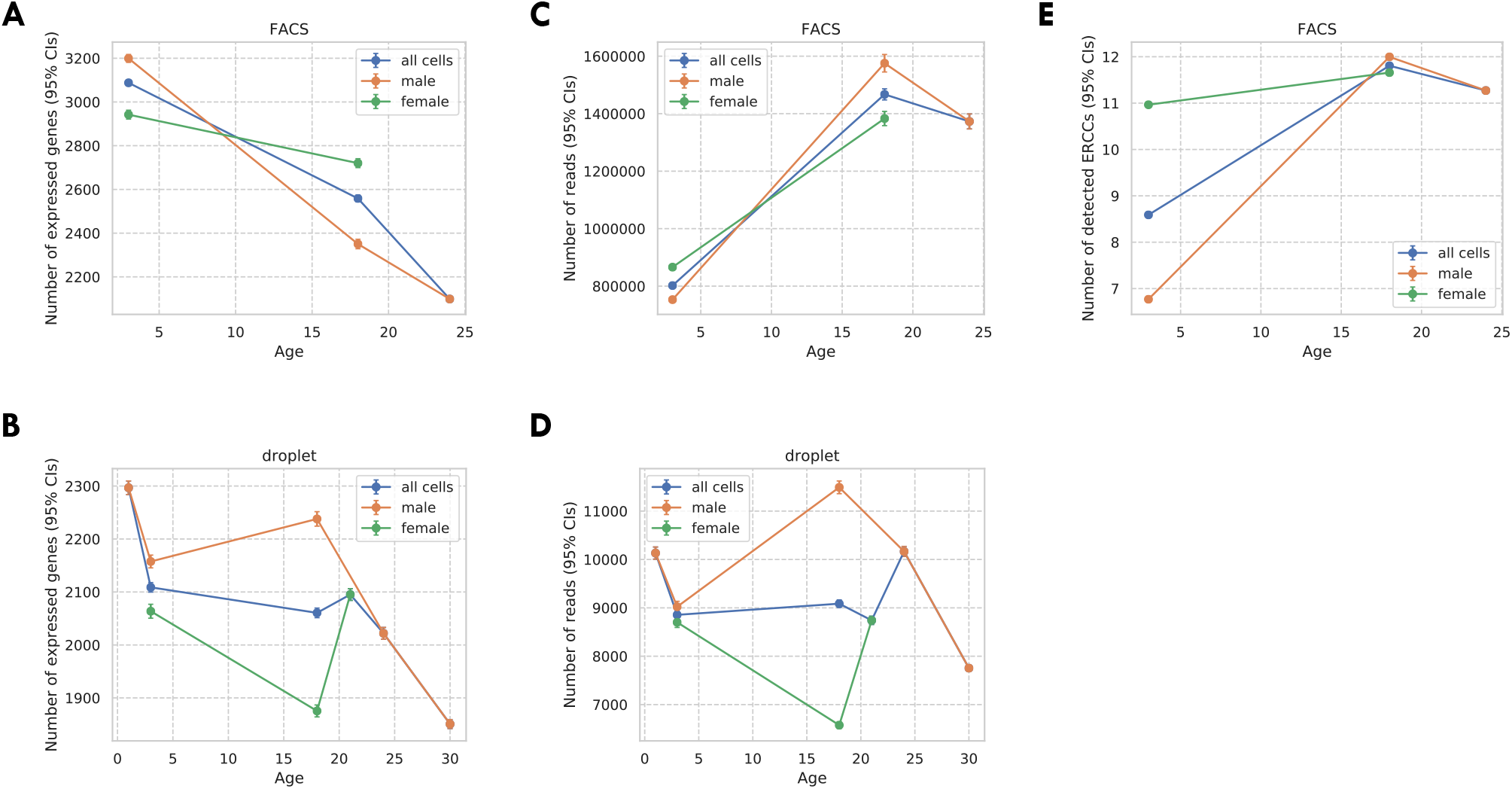
Covariates for the TMS FACS data and the TMS droplet data. A: Number of expressed genes per cell (CDR) for the TMS FACS data. B: Number of expressed genes per cell (CDR) for the TMS droplet data. C: Number of read counts per cell for the TMS FACS data. D: Number of read counts (UMIs) per cell for the TMS droplet data. E: Number of detected ERCC spike-ins per cell for the TMS FACS data. For all panels, results were presented for all cells (blue), male cells (orange), and female cells (green), separately. Also, the vertical bars represent 95% confidence intervals.

**Supplementary Figure 2.**
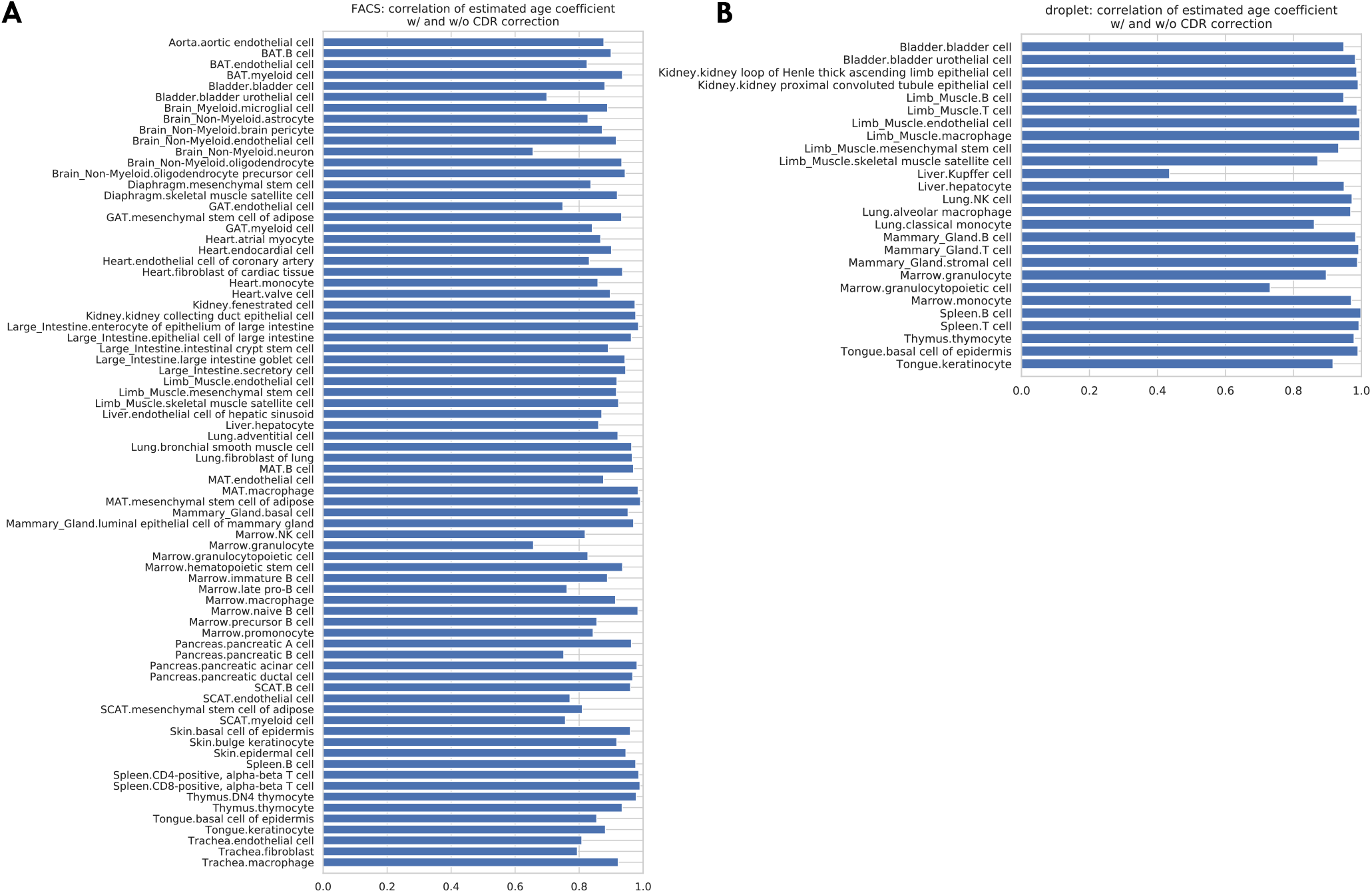
Correlations between the age coefficients estimated with and without CDR correction. The bar plots show the correlation between age coefficients estimated with and without CDR correction, across all genes, for each tissue-cell type separately. The average correlation, over all tissue-cell types, is 0.89 (std 0.08) for the FACS data (panel A) and 0.93 (std 0.12) for the droplet data (panel B).

**Supplementary Figure 3.**
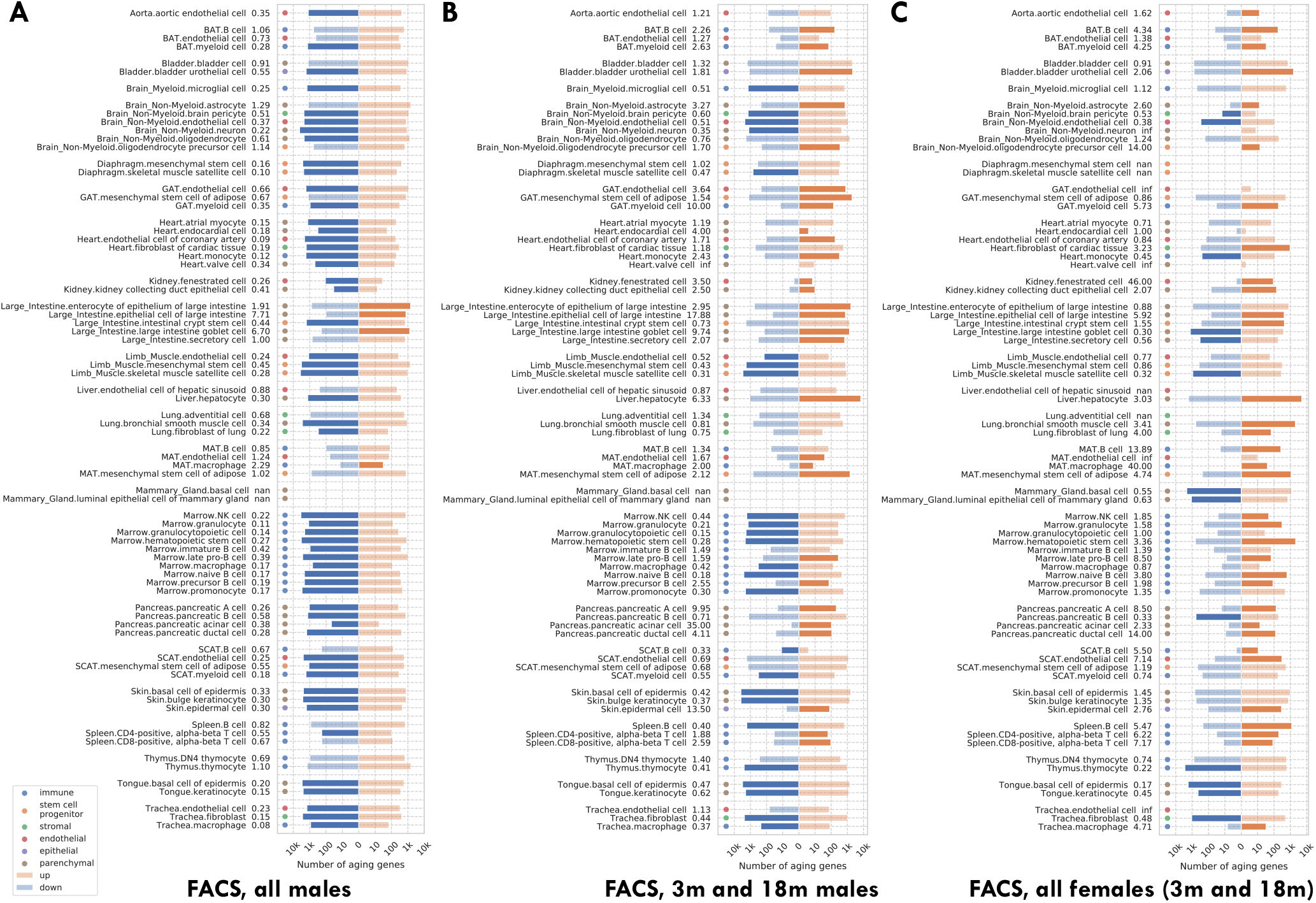
Number of significantly age-dependent genes for each tissue-cell type, from DGE analyses performed using subsets of mice in the TMS FACS data. A: Using only male mice. B: Using only 3m and 18m male mice. C: Using only female mice, which are either 3m or 18m. We can see that most tissue-cell types in panel A have significantly more down-regulated aging-related genes, but such a pattern does not exist in panels B and C. The TMS FACS data has male mice in all three time points 3m/18m/24m and female mice in only two time points 3m/18m. The comparison between panels A and B, both using male mice, indicates that the down-regulation pattern is mainly driven by the 24m mice. The comparison between panels B and C, both using 3m/18m mice, indicates that there is no systematic difference between male and female mice.

**Supplementary Figure 4.**
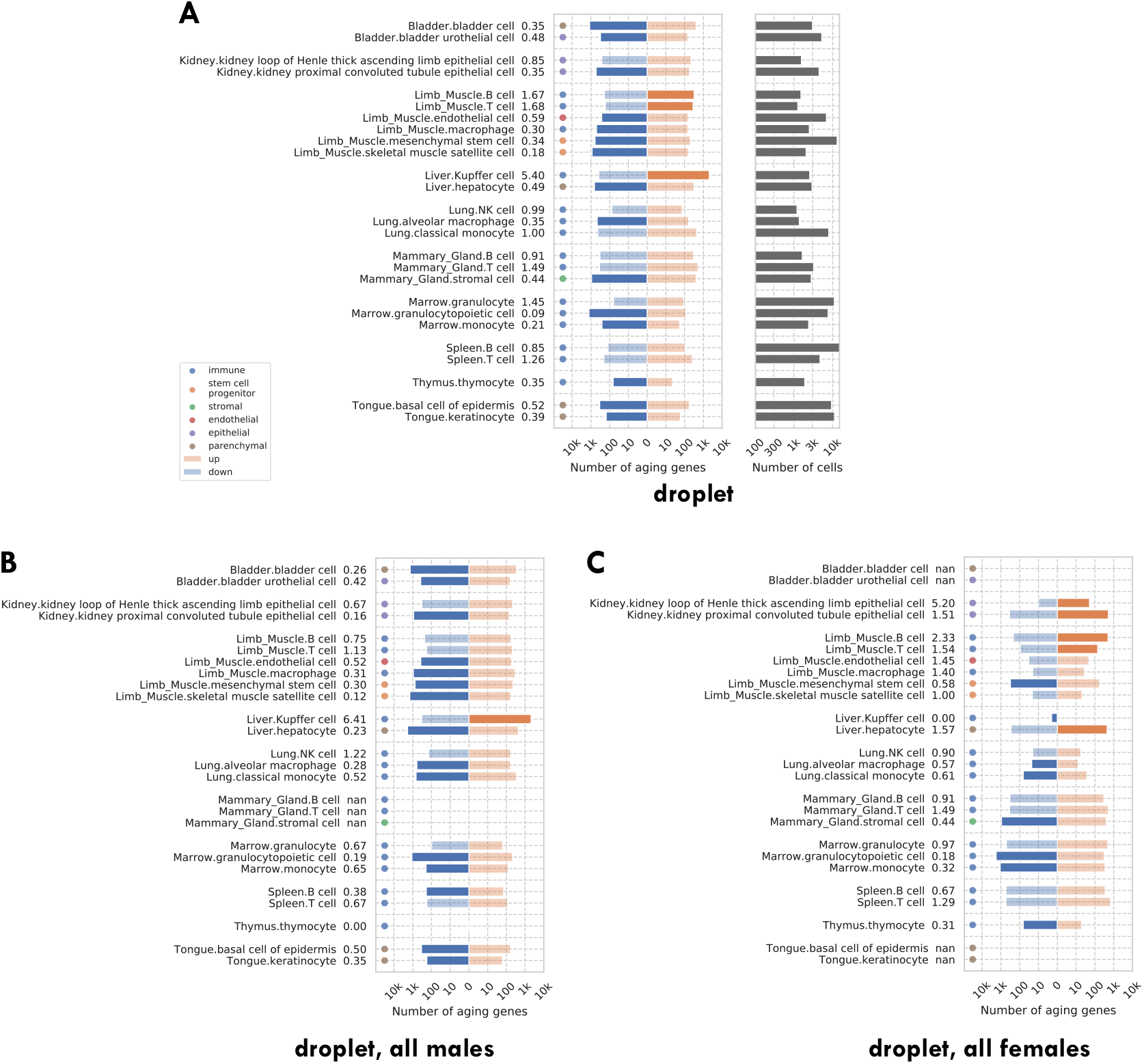
Number of significantly age-dependent genes for each tissue-cell type, from DGE analyses performed using subsets of mice in the TMS droplet data. A: Using all mice. B: Using only male mice. C: Using only female mice. We can see that there are more down-regulated aging-related genes in all three panels. The comparison between panels B and C shows that there is no systematic difference between male and female mice.

**Supplementary Figure 5.**
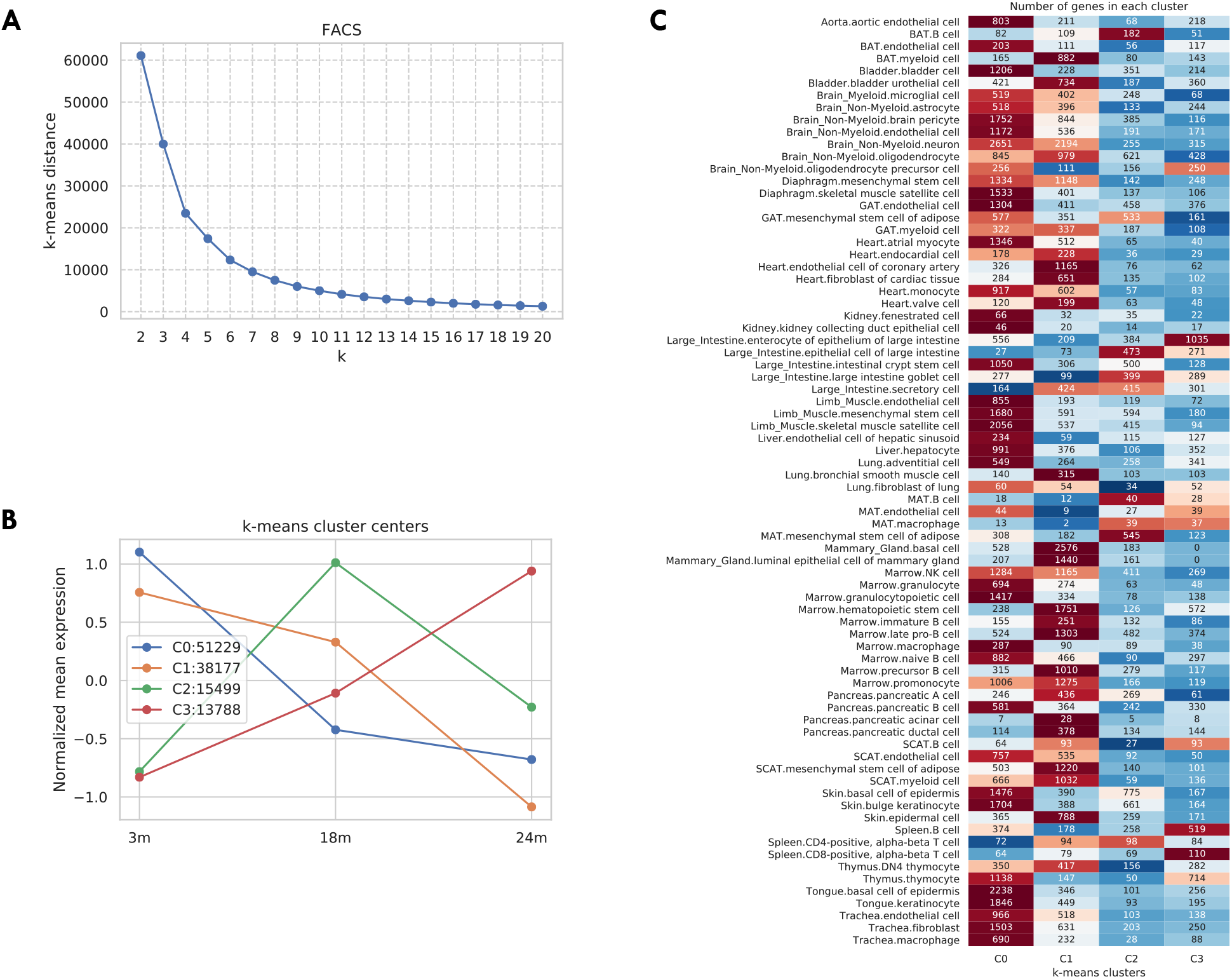
Aging trajectory of aging-related genes in the TMS FACS data. For each aging-related tissue-cell-gene tuple (defined as the gene significantly related to aging in a tissue-cell type), we computed its aging trajectory by first computing the mean expression of each time point and then z-scored the expressions over the time points (making mean 0 and std 1). Next, in order to learn the typical aging trajectories, we used the k-means clustering algorithm to cluster the aging trajectories of all those 118,693 aging-related tissue-cell-gene tuples. As shown in panel A, according to the elbow method there are k=4 clusters. We further visualized those four cluster centers in panel B. As indicated by the figure legend, most tuples are associated with cluster centers 0 and 1, corresponding to monotonically decreasing gene expressions from 3m to 24m. Interestingly, 15,499 tuples are associated with cluster center 2 whose gene expressions first increased from 3m to 18m and then decreased from 18m to 24m. Also, 13,788 tuples are associated with cluster center 3, corresponding to a monotonically increasing aging trajectory. In panel C, we further counted those tuples by tissue-cell types. While most tissue-cell types have aging-related genes belonging to cluster centers 0 and 1 corresponding to monotonically decreasing trajectories, B cells in the brown adipose tissue (BAT), epithelial cells in the large intestine, and mesenchymal stem cells of adipose in the mesenchymal adipose tissue (MAT) have more aging-related genes in cluster 2, where the gene expressions increased from 3m to 18m and then decreased from 18m to 24m. In addition, cell types in the MAT, large intestine, and spleen have more up-regulated aging-related genes that correspond to clusters 2 and 3, consistent with Fig. 1B.

**Supplementary Figure 6.**
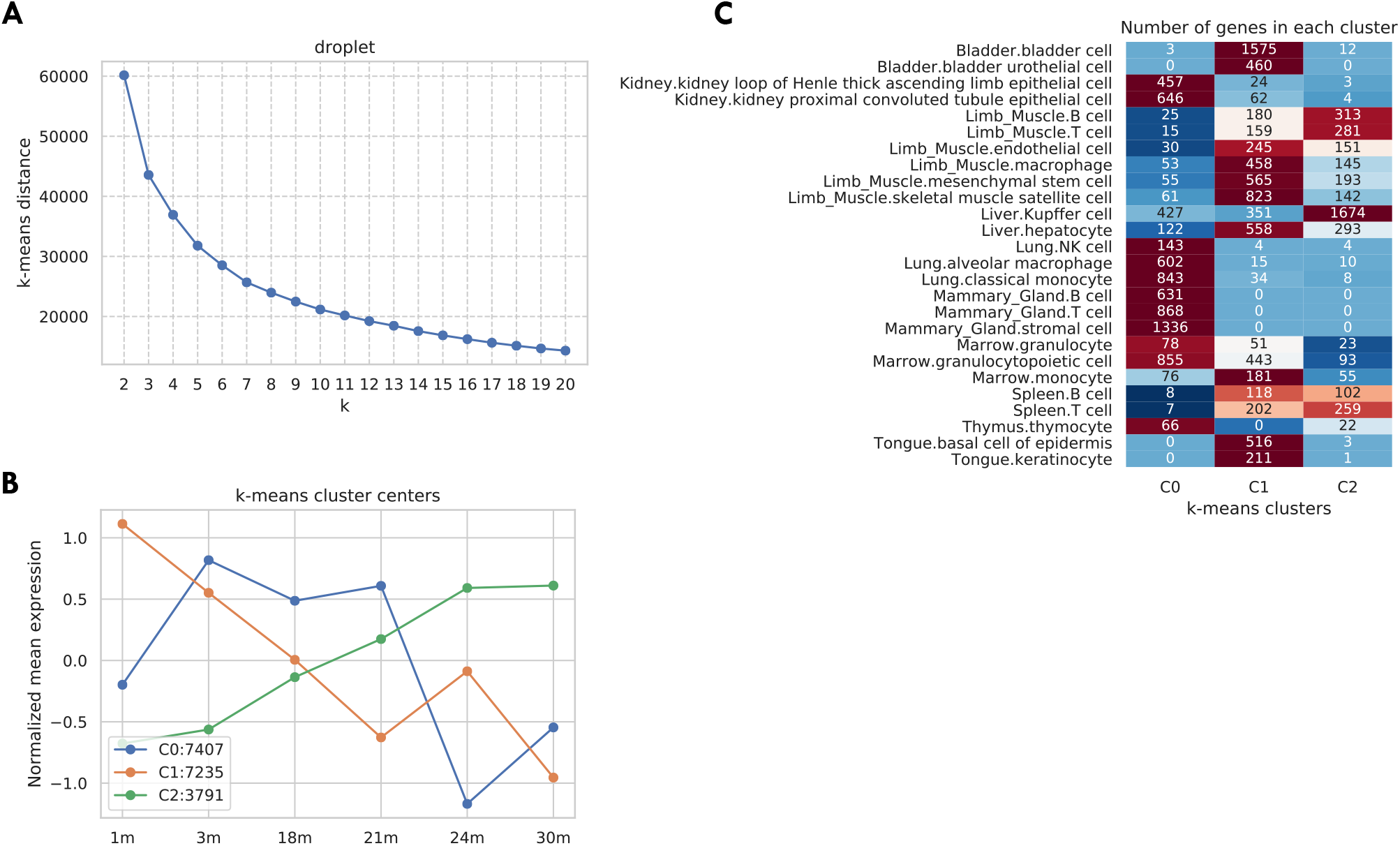
Aging trajectory of aging-related genes in the TMS droplet data. We computed and clustered the aging trajectories for genes in the TMS droplet data similar to Supp. Fig. 5. As shown in panel A, according to the elbow method there are k=3 clusters. We further visualized those three cluster centers in panel B. As indicated by the figure legend, most tuples are associated with cluster centers 0 and 1, corresponding to a general decreasing pattern from 1m to 30m. 3,791 tuples are associated with cluster center 2, corresponding to a monotonically increasing aging trajectory. In panel C, we further counted those tuples by tissue-cell types. While most tissue-cell types have aging-related genes belonging to clusters 0 and 1 corresponding to generally decreasing trajectories, cell types in the limb muscle, and Kupffer cells in the kidney have more up-regulated aging-related genes that correspond to cluster 2, consistent with Supp. Fig. 4A.

**Supplementary Figure 7.**
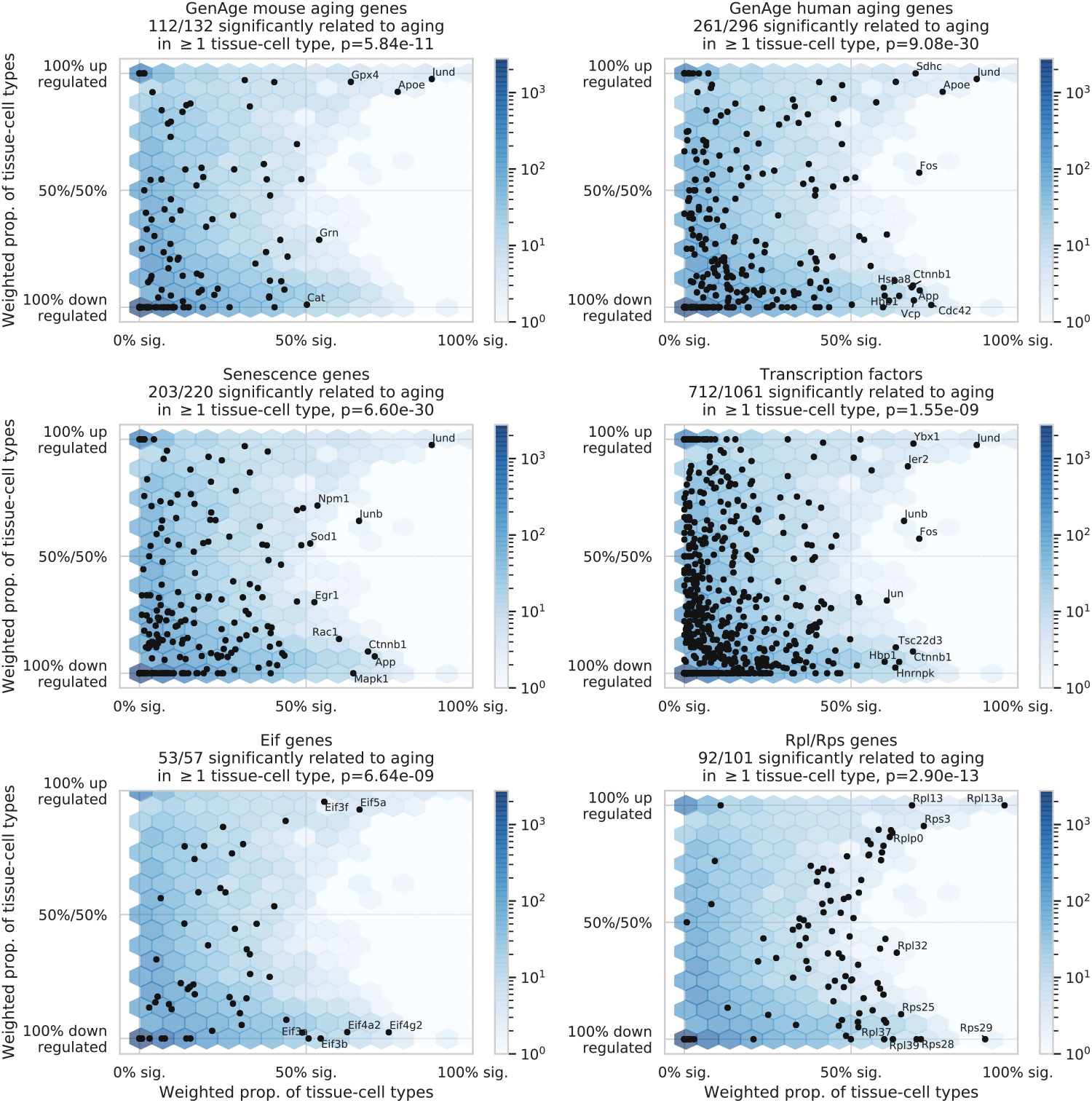
Comparison between aging-related genes discovered in the TMS FACS data and other gene sets, including the mouse aging genes and human aging genes in the GenAge database^5^, senescence genes as provided by the IPA software^3^, transcription factors, eukaryotic initiation factors, and ribosomal protein genes (Rpl/Rps genes). The figure axes are the same as Figure 2A. We observe significant overlap between the FACS aging-related genes and all six gene sets, as quantified by the Fisher’s exact tests with p-values provided in the panel titles.

**Supplementary Figure 8.**
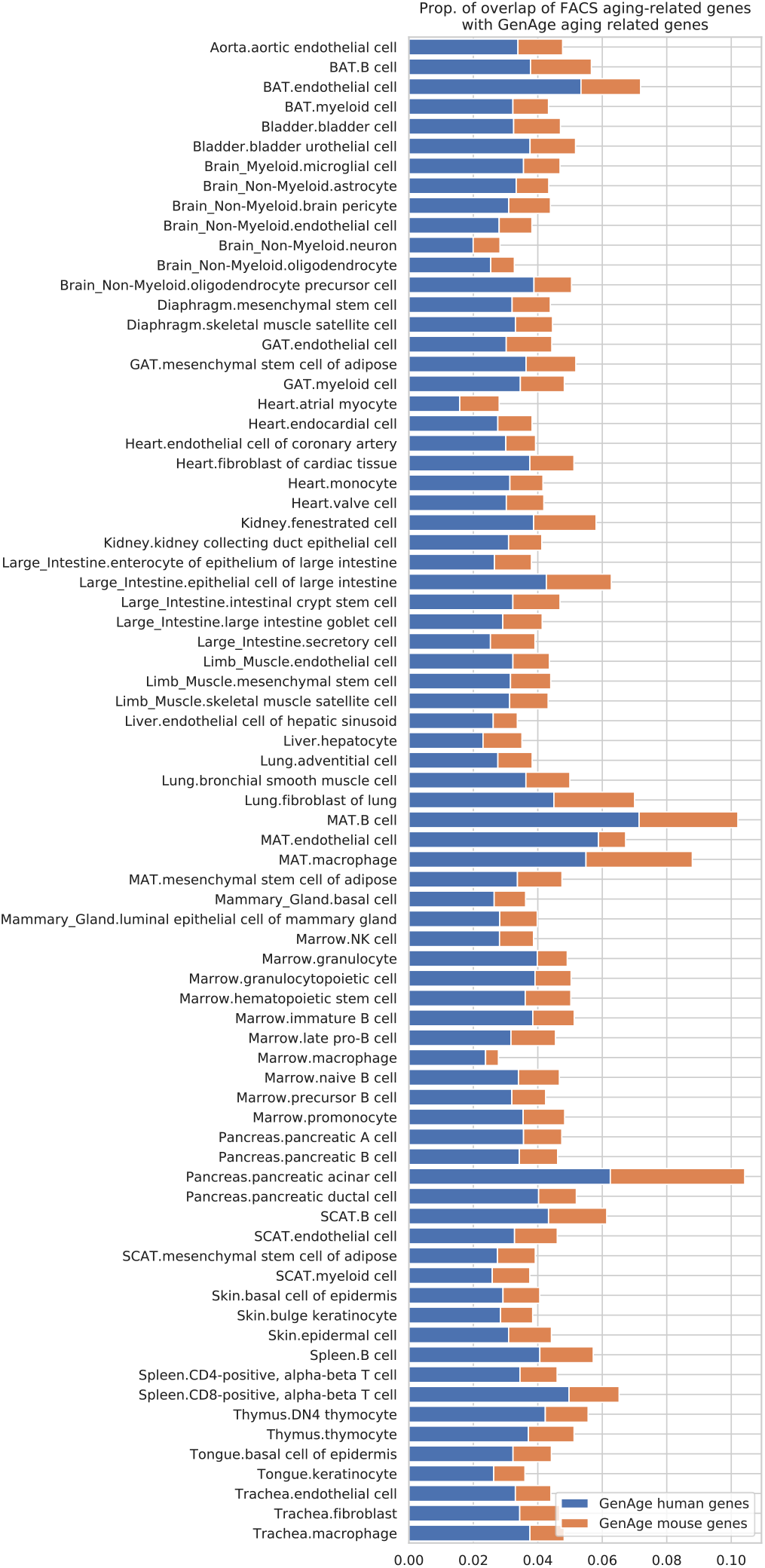
Overlap between the TMS FACS aging-related genes and the GenAge aging markers. The figure shows, for each tissue-cell type, the proportion of aging-related genes that are also shared by the GenAge human aging genes (blue) and the GenAge mouse aging genes (orange). No tissue-cell type is particularly enriched with the GenAge human aging genes. There are 303 human aging genes and 136 mouse aging genes in the GenAge database (97 genes shared by the two sets), explaining why the FACS aging-related genes overlap more with the GenAge human aging genes.

**Supplementary Figure 9.**
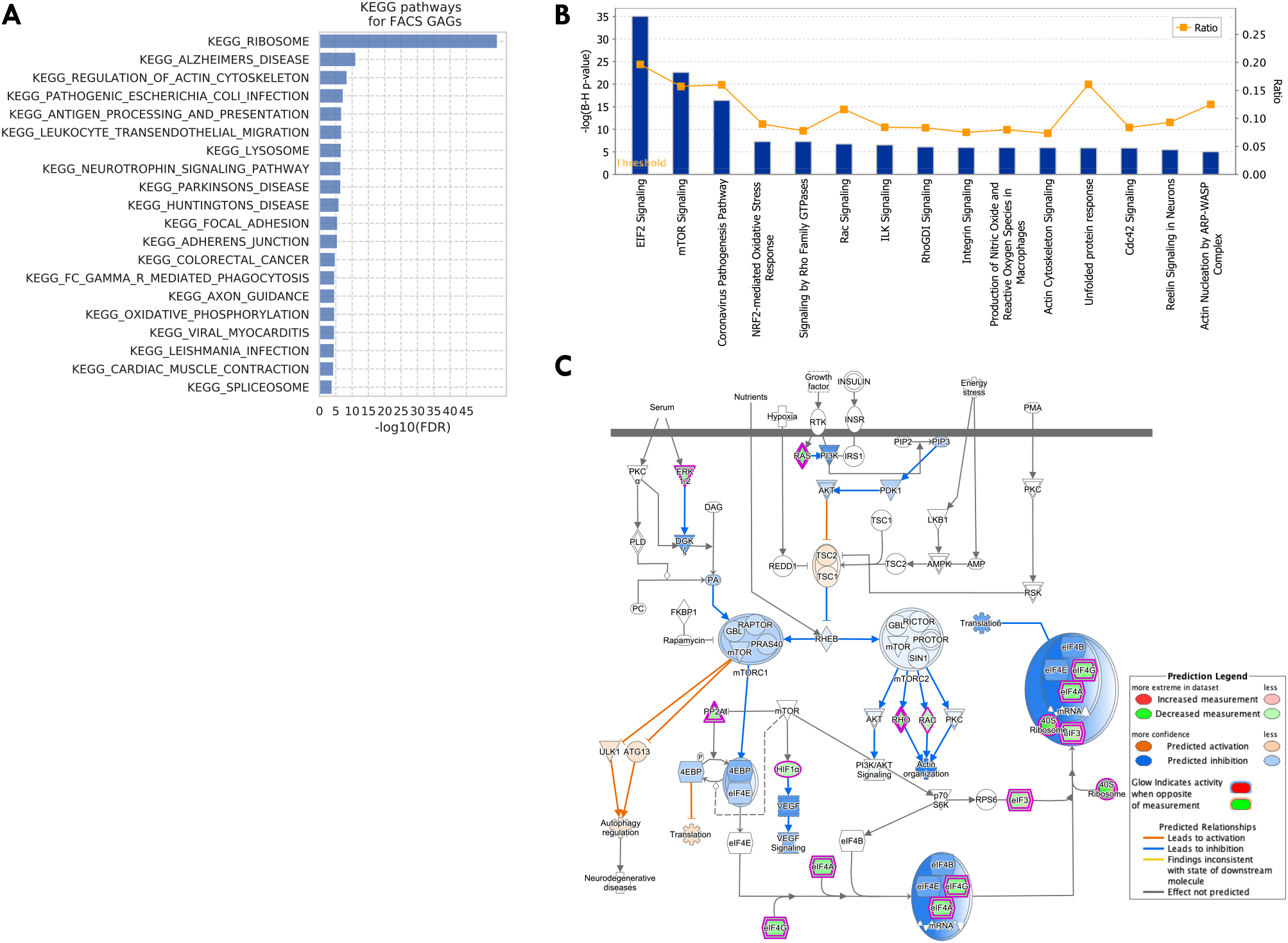
KEGG pathways and IPA pathways for the 330 TMS FACS GAGs. A: Top 20 KEGG pathways for the GAGs. B: Top IPA pathways for the GAGs. The ratio represents the proportion of pathway genes that are also GAGs. C: Regulation prediction for mTOR pathway by IPA. Overall, the pathway is predicted to be down-regulated, with related processes including translation, actin organization, and VEGF (vascular endothelial growth factor) signaling.

**Supplementary Figure 10.**
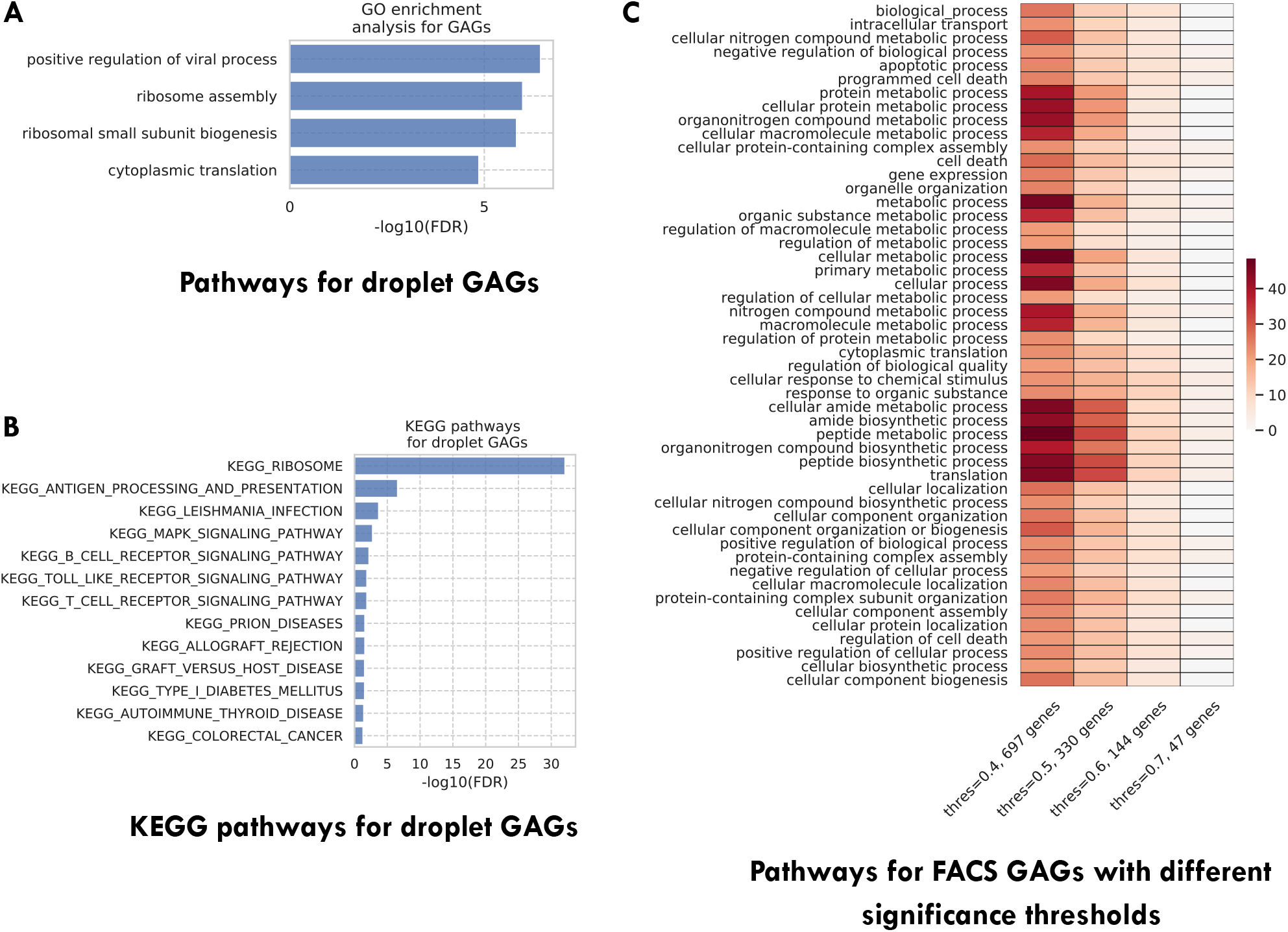
Additional pathway enrichment analysis results. A: Top GO biological pathways for the TMS droplet GAGs. B: Top 20 KEGG pathways for the TMS droplet GAGs. C: Top GO biological pathways for the TMS FACS GAGs selected with different significance thresholds (as indicated in the x-axis). The color represents the negative log10 FDR. As we can see, GAGs selected using different thresholds are associated with similar pathways.

**Supplementary Figure 11.**
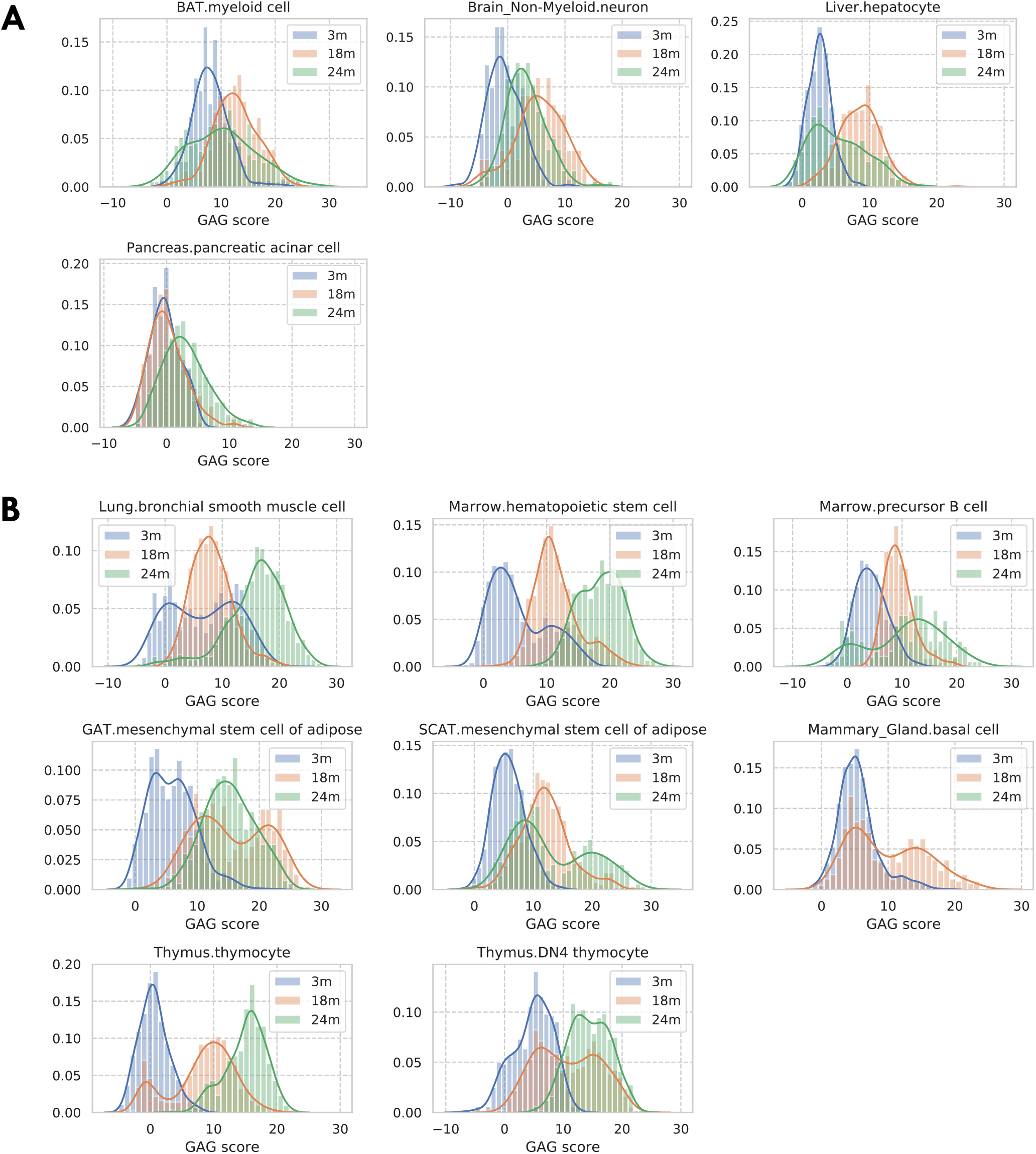
Distribution of GAG scores across cells; stratified by tissue-cell types and grouped by mice’s chronological age. Most tissue-cell types have lower GAG scores for young cells and higher GAG scores for old cells. However, as shown by the GAG score distribution plots in panel A, we found four tissue-cell types whose GAG scores are similar between age groups, suggesting that the cells from these four tissue-cell types do not change much during aging. We also found eight tissue-cell types that each has a subpopulation of cells whose GAG scores are more similar to that of cells from a different age group (panel B). Specifically, bronchial smooth muscle cells in the lung and hematopoietic stem cells in the marrow have a subpopulation of 3m cells that has larger GAG scores like old cells, suggesting that those young cells may have an aging status similar to that of old cells. On the other hand, each of the other six tissue-cell types has a subpopulation of old cells that has much smaller GAG scores, implying that those old cells may have an aging status similar to that of young cells.

**Supplementary Figure 12.**
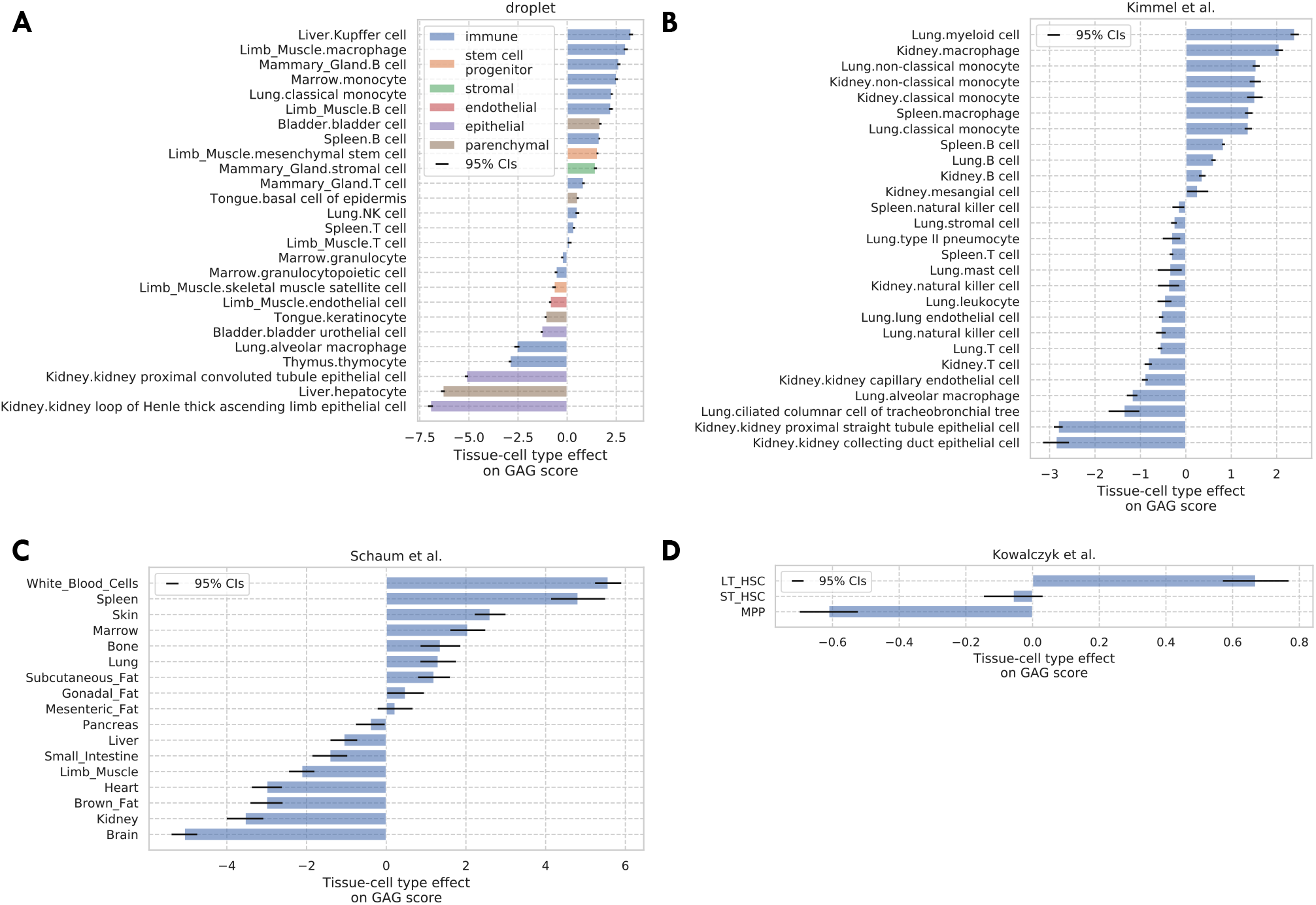
Tissue-cell GAG score effects for the four validation datasets. A: TMS droplet data. B: Kimmel et al. data^1^. C: The bulk data^4^. D: Kowalczyk et al. data^2^.

**Supplementary Figure 13.**
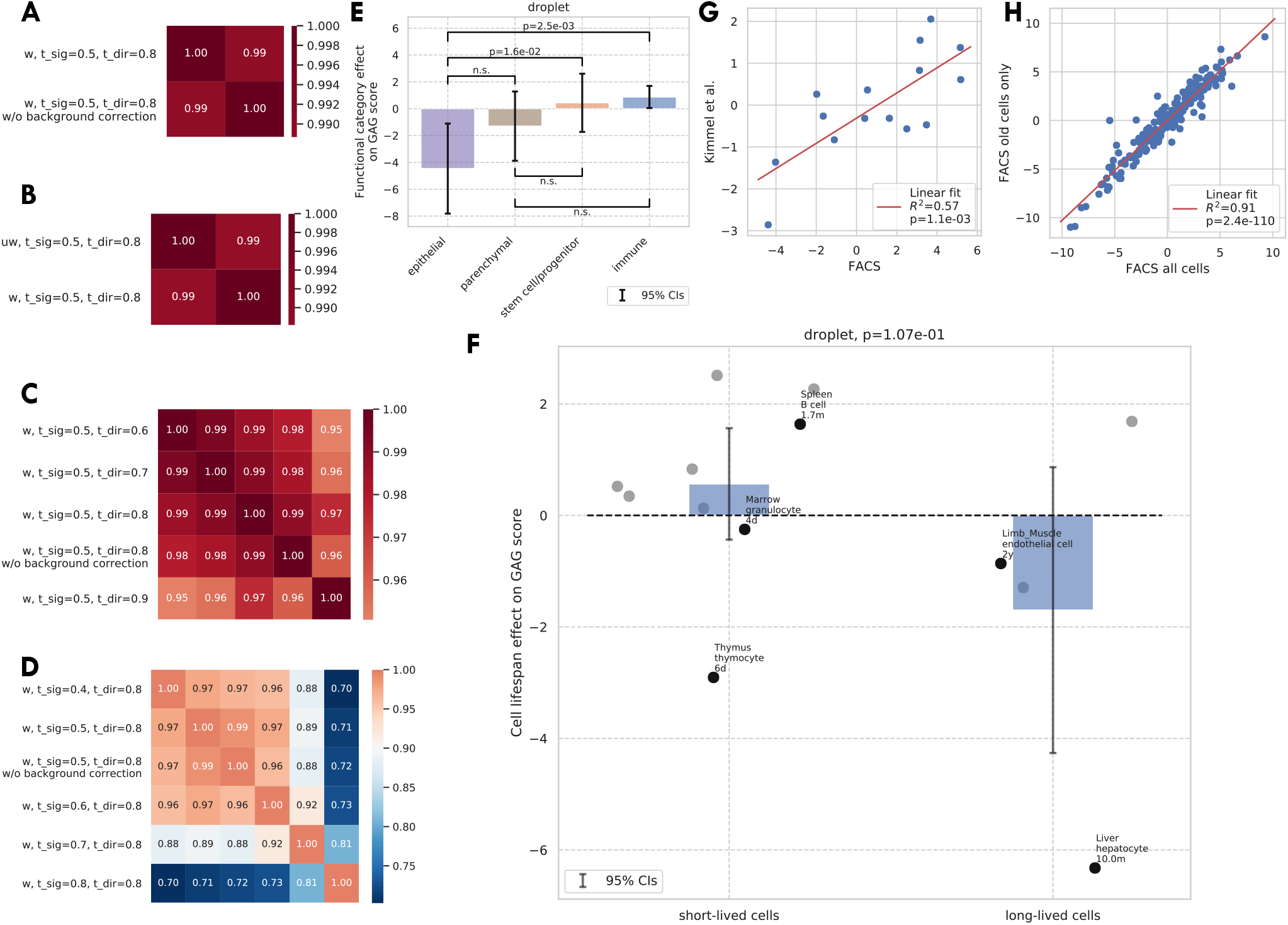
Additional validations of the GAG score. A-D: Correlation between the GAG scores computed using different scoring rules. The original GAG score was computed using genes that are significantly related to aging in >50% of weighted tissue-cell types and have the same directionality in >80% of weighted tissue-cell types (i.e., “w, t_sig=0.5, t_dir=0.8”). After that, a cell-wise background correction was performed. Panel A shows that the cell-wise background correction does not significantly change the GAG score. Panel B shows that weighted and unweighted tissue-cell types (w/uw) give similar results. Panel C shows that different directionality thresholds (t_dir) give similar results. Panel D shows that different significance thresholds (t_sig) give similar results, except that when t_sig=0.9, which is probably too stringent. E-F: Effects of functional categories (panel D) and binary cell lifespan (panel E) on the GAG score in the TMS droplet data, meta-analyzed over all tissue-cell types in the group. A positive y-value means that the cells in the group (functional category group for panel E and binary lifespan group for panel F) have higher GAG score values than other cells of the same age and sex. 95% confidence intervals and nominal p-values are provided to quantify the differences between categories. In panel F, the average cell lifespan annotations are also provided for a subset of tissue-cell types. G: The comparison between the tissue-cell GAG score effects estimated from the FACS data (x-axis) and from the Kimmel et al. data^1^ (y-axis). Each dot corresponds to a tissue-cell type, and a linear fit is provided, showing that the estimates are consistent. H: The comparison between the tissue-cell GAG score effects estimated using all cells in the FACS data (x-axis) and using only old cells in the FACS data (y-axis). Each dot corresponds to a tissue-cell type, and a linear fit is provided, showing that the estimates are consistent.

**Supplementary Figure 14.**
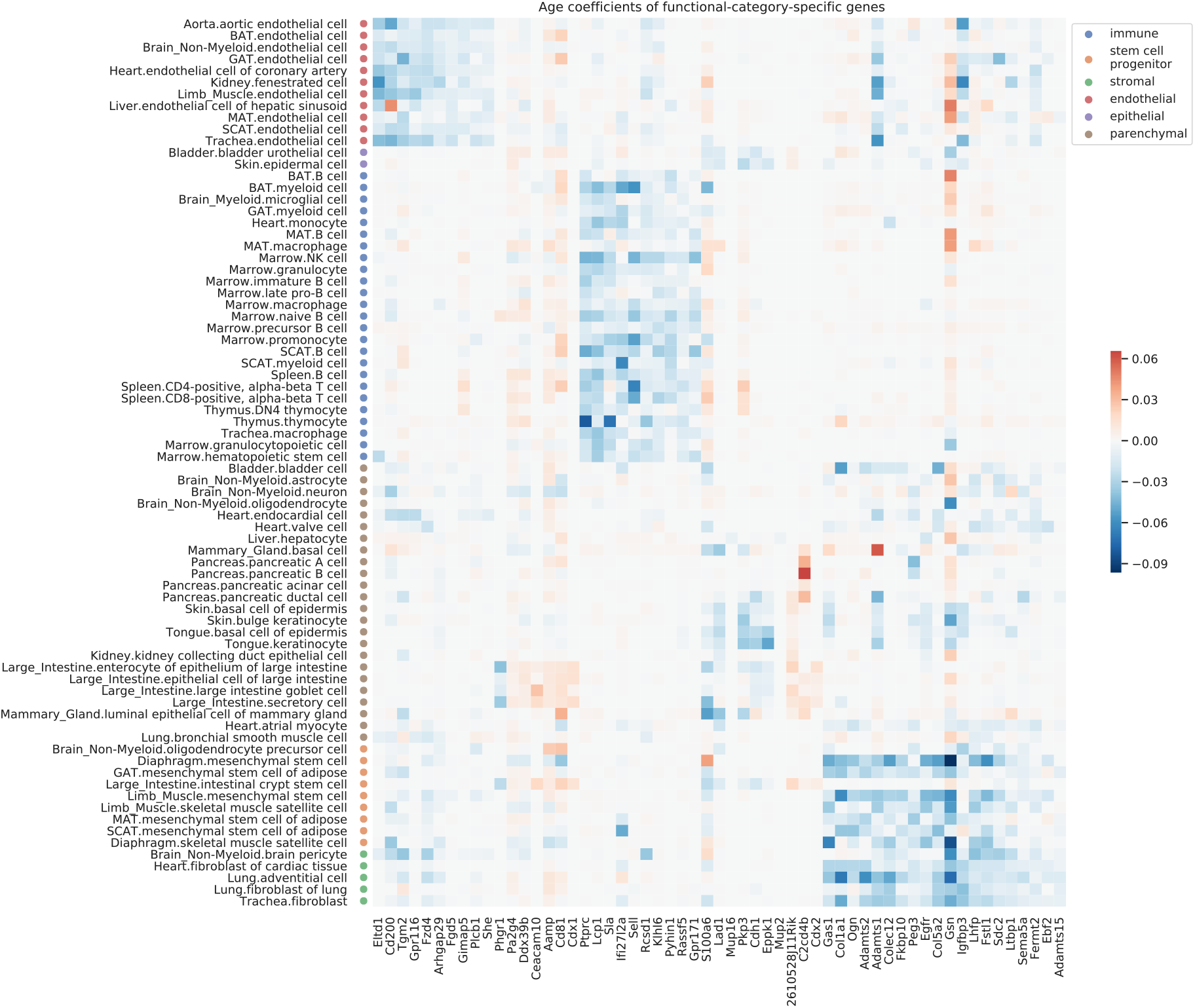
Age coefficients of top functional-category-specific genes. Top 10 genes with the most significant p-values were selected for each category, ordered from left to right as the endothelial, epithelial, immune, stem/progenitor, stromal, and parenchymal category. For example, the upper-left block corresponds to genes specific to endothelial cells. The color represents the age coefficients, in the unit of log10 fold change per month.

**Supplementary Figure 15.**
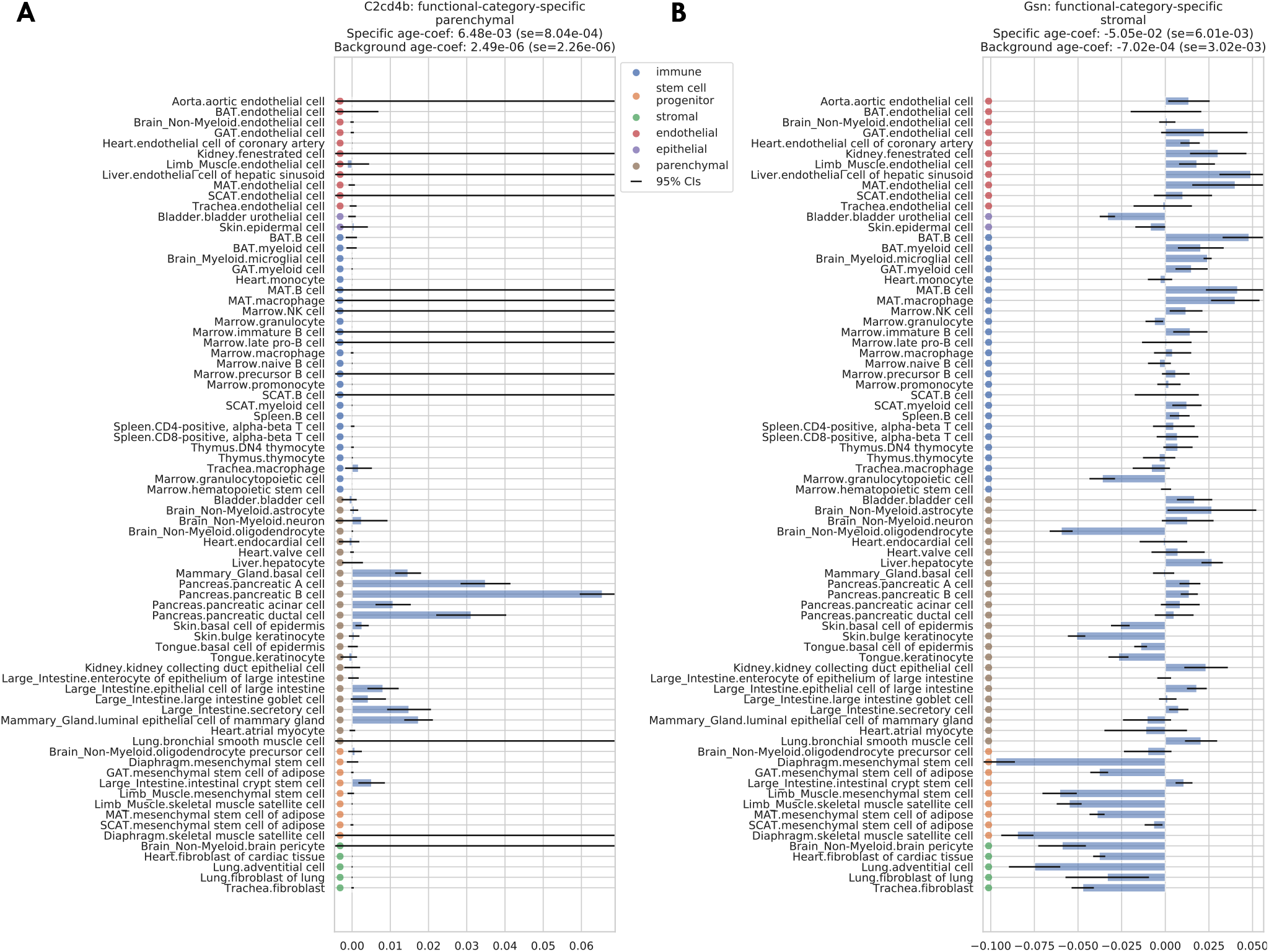
Two examples of functional-category-specific genes. A: *C2cd4b* which is parenchymal-specific. B: *Gsn* which is stromal-specific. Estimated age coefficients are provided for each tissue-cell type with 95% confidence intervals. Within-set and outside-set meta age coefficients, estimated via meta-analysis, are provided in the panel titles.

**Supplementary Figure 16.**
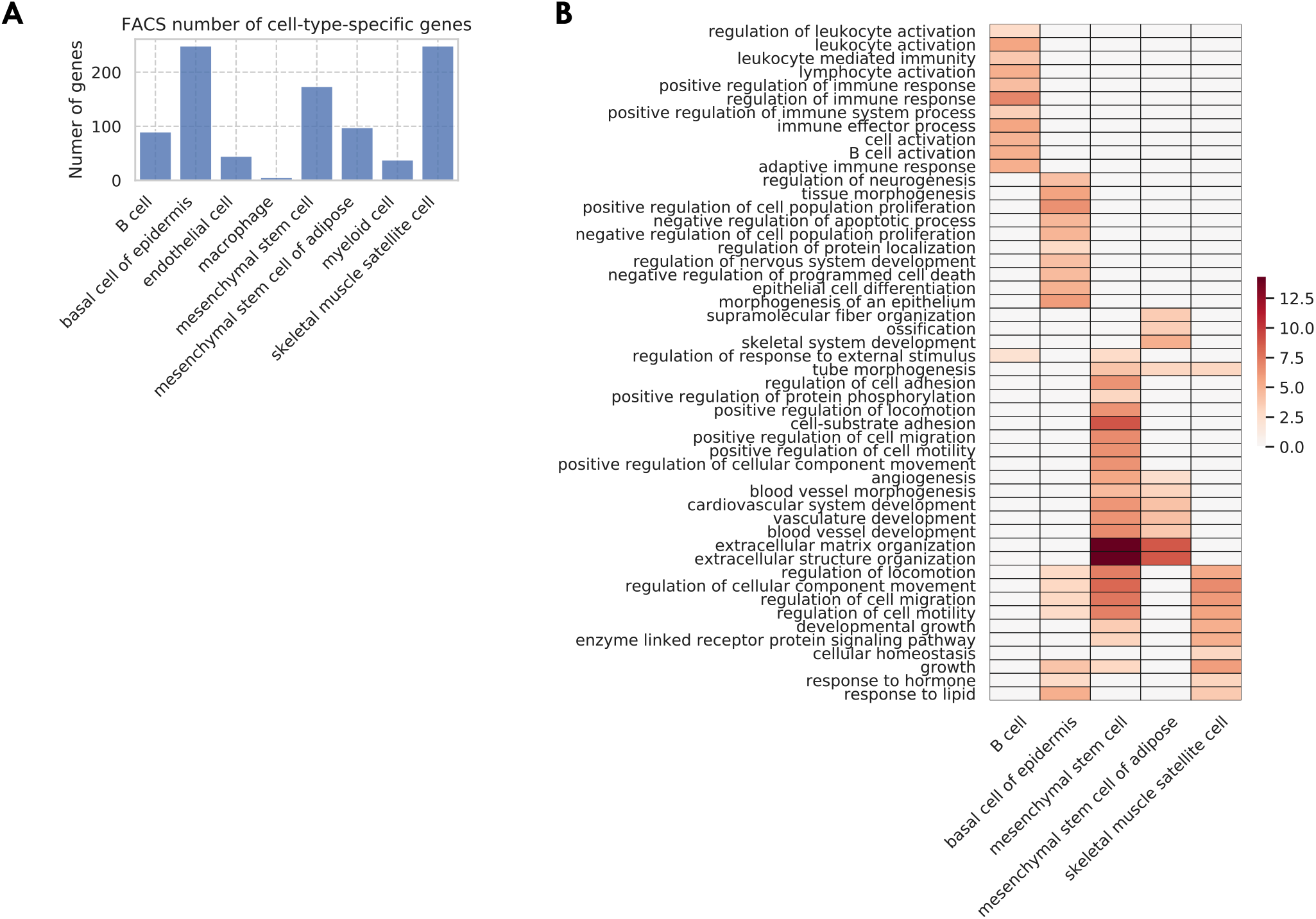
Cell-type-specific genes. A: Number of cell-type-specific genes. B: GO biological pathways for the cell-type-specific genes, with color representing the negative log10 FDR.

**Supplementary Figure 17.**
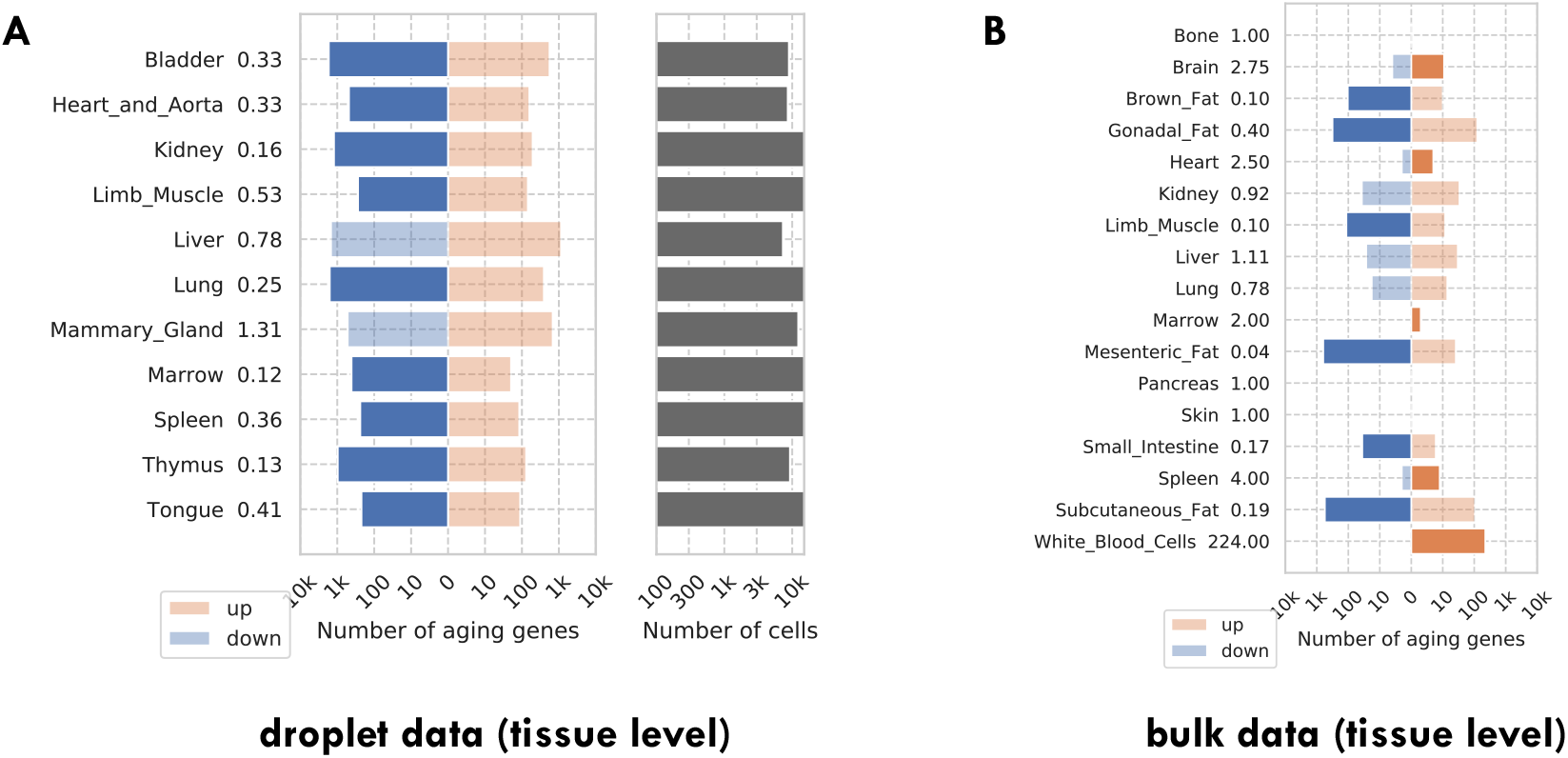
Number of discoveries for each tissue. A: TMS droplet data. B: The bulk data. The left panels show the number of aging genes (discoveries) for each tissue, broken down into the number of up-regulated genes (orange), and the number of down-regulated genes (blue), with the numbers on the left showing the ratio (up/down). Tissue types with significantly more up-/down- regulated genes (ratio>1.5) are highlighted in the solid color. The right panels show the number of cells for each tissue.

**Supplementary Figure 18.**
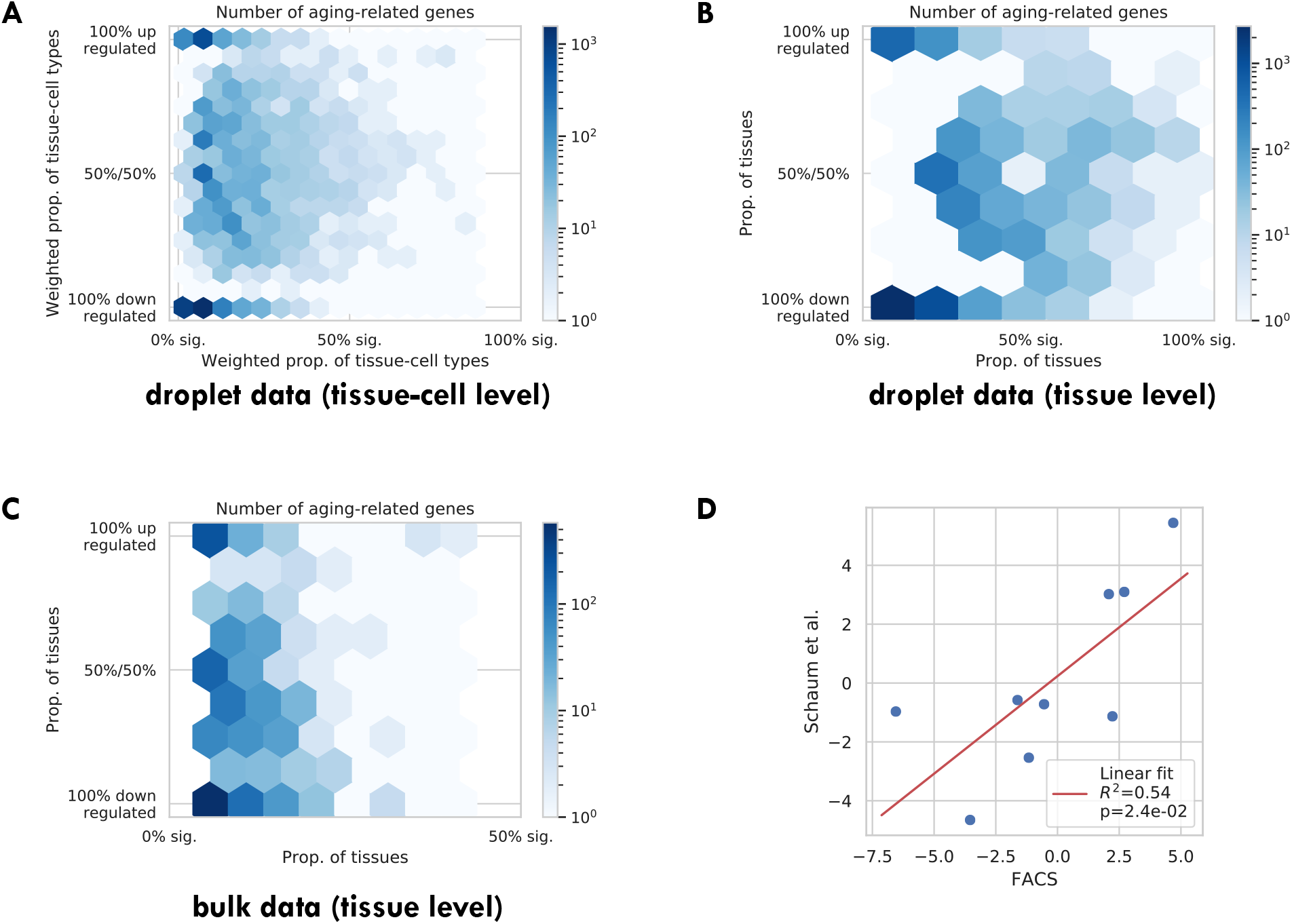
Tissue-cell validations. A: Tissue-cell level aging-related genes for the TMS droplet data with color indicating the number of genes. B: Tissue-level aging-related genes for the TMS droplet data with color indicating the number of genes. C: Tissue-level aging-related genes for the bulk data with color indicating the number of genes. D: Comparison between the FACS tissue-level GAG score effects and the bulk data tissue-level GAG score effects. Each dot corresponds to a tissue type, and a linear fit is provided showing that the estimates are consistent.

**Supplementary Figure 19.**
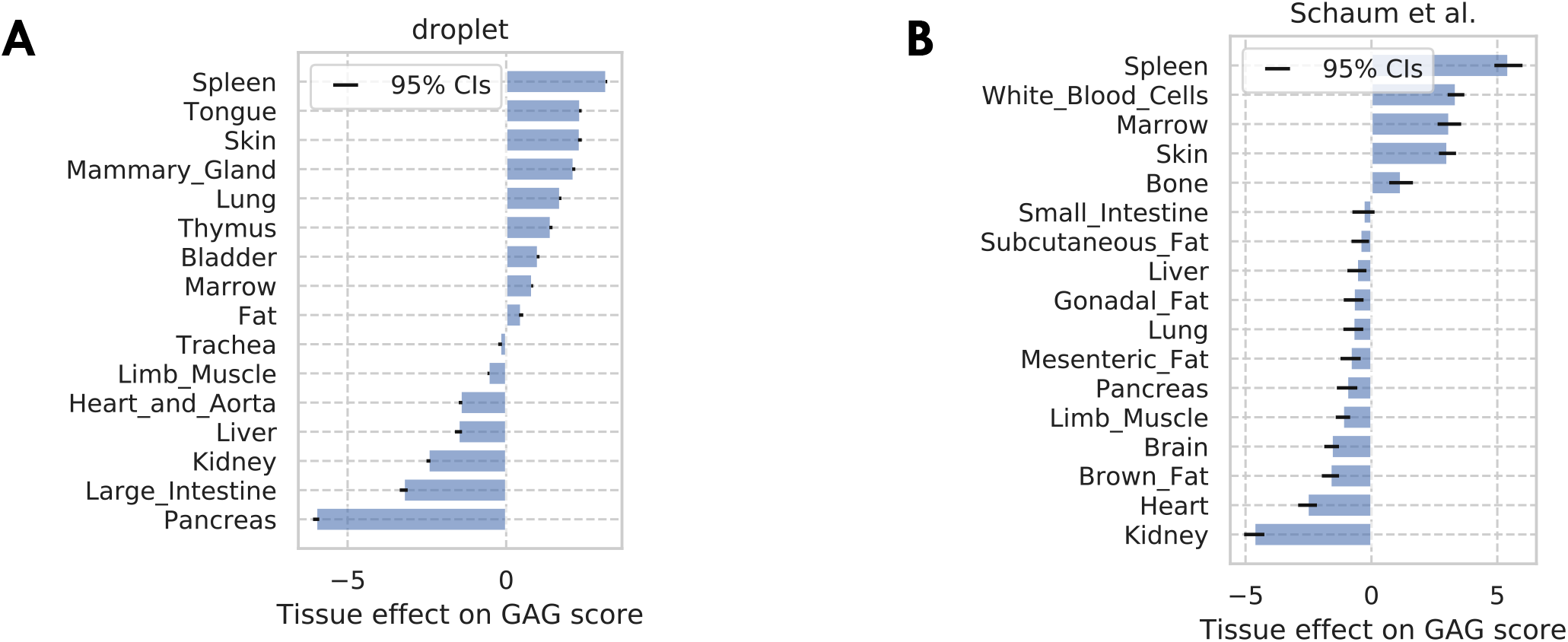
Tissue-level GAG score effects. A: TMS droplet data. B: The bulk data. The results are consistent with the result in the FACS data as shown in Fig. 5C, with the immune tissues (spleen, white blood cells, marrow, thymus) being at the top, and pancreas being at the bottom.

## References

1. Aleksandra S Anisimova, Alexander I Alexandrov, Nadezhda E Makarova, Vadim N Gladyshev, and Sergey E Dmitriev. Protein synthesis and quality control in aging. Aging (Albany NY), 10(12):4269, 2018.

2. Nir Barzilai, Derek M Huffman, Radhika H Muzumdar, and Andrzej Bartke. The critical role of metabolic pathways in aging. Diabetes, 61(6):1315–1322, 2012.

3. Yoav Benjamini and Yosef Hochberg. Controlling the false discovery rate: a practical and powerful approach to multiple testing. Journal of the Royal statistical society: series B (Methodological), 57(1):289–300, 1995.

4. Olaf Bergmann, Sofia Zdunek, Anastasia Felker, Mehran Salehpour, Kanar Alkass, Samuel Bernard, Staffan L Sjostrom, Mirosława Szewczykowska, Teresa Jackowska, Cris Dos Remedios, et al. Dynamics of cell generation and turnover in the human heart. Cell, 161(7):1566–1575, 2015.

5. Judith Campisi. Aging, cellular senescence, and cancer. Annual review of physiology, 75:685–705, 2013.

6. Samuel H Cheshier, Sean J Morrison, Xinsheng Liao, and Irving L Weissman. In vivo proliferation and cell cycle kinetics of long-term self-renewing hematopoietic stem cells. Proceedings of the National Academy of Sciences, 96(6):3120–3125, 1999.

7. Adam S Darwich, Umair Aslam, Darren M Ashcroft, and Amin Rostami-Hodjegan. Meta-analysis of the turnover of intestinal epithelia in preclinical animal species and humans. Drug Metabolism and Disposition, 42(12):2016–2022, 2014.

8. David DeTomaso, Matthew G Jones, Meena Subramaniam, Tal Ashuach, J Ye Chun, and Nir Yosef. Functional interpretation of single cell similarity maps. Nature communications, 10(1):1–11, 2019.

9. Rafael Arrojo e Drigo, Varda Lev-Ram, Swati Tyagi, Ranjan Ramachandra, Thomas Deerinck, Eric Bushong, Sebastien Phan, Victoria Orphan, Claude Lechene, Mark H Ellisman, et al. Age mosaicism across multiple scales in adult tissues. Cell metabolism, 30(2):343–351, 2019.

10. Maria Ermolaeva, Francesco Neri, Alessandro Ori, and K Lenhard Rudolph. Cellular and epigenetic drivers of stem cell ageing. Nature Reviews Molecular Cell Biology, 19(9):594, 2018.

11. Greg Finak, Andrew McDavid, Masanao Yajima, Jingyuan Deng, Vivian Gersuk, Alex K Shalek, Chloe K Slichter, Hannah W Miller, M Juliana McElrath, Martin Prlic, et al. Mast: a flexible statistical framework for assessing transcriptional changes and characterizing heterogeneity in single-cell rna sequencing data. Genome biology, 16(1):278, 2015.

12. Jason G Fleischer, Roberta Schulte, Hsiao H Tsai, Swati Tyagi, Arkaitz Ibarra, Maxim N Shokhirev, Ling Huang, Martin W Hetzer, and Saket Navlakha. Predicting age from the transcriptome of human dermal fibroblasts. Genome biology, 19(1):221, 2018.

13. DA Fulcher and A Basten. B cell life span: a review. Immunology and cell biology, 75(5):446–455, 1997.

14. Barbara Geering, Christina Stoeckle, Sébastien Conus, and Hans-Uwe Simon. Living and dying for inflammation: neutrophils, eosinophils, basophils. Trends in immunology, 34(8):398–409, 2013.

15. Martin Guilliams, Alexander Mildner, and Simon Yona. Developmental and functional heterogeneity of monocytes. Immunity, 49(4):595–613, 2018.

16. Gunsagar S. Gulati, Shaheen S. Sikandar, Daniel J. Wesche, Anoop Manjunath, Anjan Bharadwaj, Mark J. Berger, Francisco Ilagan, Angera H. Kuo, Robert W. Hsieh, Shang Cai, Maider Zabala, Ferenc A. Scheeren, Neethan A. Lobo, Dalong Qian, Feiqiao B. Yu, Frederick M. Dirbas, Michael F. Clarke, and Aaron M. Newman. Single-cell transcriptional diversity is a hallmark of developmental potential. Science, 367(6476):405–411, 2020.

17. Lorna W Harries, Dena Hernandez, William Henley, Andrew R Wood, Alice C Holly, Rachel M Bradley-Smith, Hanieh Yaghootkar, Ambarish Dutta, Anna Murray, Timothy M Frayling, et al. Human aging is characterized by focused changes in gene expression and deregulation of alternative splicing. Aging cell, 10(5):868–878, 2011.

18. B Hobson and J Denekamp. Endothelial proliferation in tumours and normal tissues: continuous labelling studies. British journal of cancer, 49(4):405–413, 1984.

19. Alice C Holly, David Melzer, Luke C Pilling, William Henley, Dena G Hernandez, Andrew B Singleton, Stefania Bandinelli, Jack M Guralnik, Luigi Ferrucci, and Lorna W Harries. Towards a gene expression biomarker set for human biological age. Aging cell, 12(2):324–326, 2013.

20. Steve Horvath. Dna methylation age of human tissues and cell types. Genome biology, 14(10):3156, 2013.

21. Simon C Johnson, Peter S Rabinovitch, and Matt Kaeberlein. mtor is a key modulator of ageing and age-related disease. Nature, 493(7432):338–345, 2013.

22. Juulia Jylhävä, Nancy L Pedersen, and Sara Hägg. Biological age predictors. EBioMedicine, 21:29–36, 2017.

23. Jacob C Kimmel, Lolita Penland, Nimrod D Rubinstein, David G Hendrickson, David R Kelley, and Adam Z Rosenthal. Murine single-cell rna-seq reveals cell-identity-and tissue-specific trajectories of aging. Genome research, 29(12):2088–2103, 2019.

24. Maranke I Koster. Making an epidermis. Annals of the New York Academy of Sciences, 1170:7, 2009.

25. Monika S Kowalczyk, Itay Tirosh, Dirk Heckl, Tata Nageswara Rao, Atray Dixit, Brian J Haas, Rebekka K Schneider, Amy J Wagers, Benjamin L Ebert, and Aviv Regev. Single-cell rna-seq reveals changes in cell cycle and differentiation programs upon aging of hematopoietic stem cells. Genome research, 25(12):1860–1872, 2015.

26. Andreas Krämer, Jeff Green, Jack Pollard Jr, and Stuart Tugendreich. Causal analysis approaches in ingenuity pathway analysis. Bioinformatics, 30(4):523–530, 2013.

27. Ina Kycia, Brooke N Wolford, Jeroen R Huyghe, Christian Fuchsberger, Swarooparani Vadlamudi, Romy Kursawe, Ryan P Welch, Ricardo d’Oliveira Albanus, Asli Uyar, Shubham Khetan, et al. A common type 2 diabetes risk variant potentiates activity of an evolutionarily conserved islet stretch enhancer and increases c2cd4a and c2cd4b expression. The American Journal of Human Genetics, 102(4):620–635, 2018.

28. Laurent Lamalice, Fabrice Le Boeuf, and Jacques Huot. Endothelial cell migration during angiogenesis. Circulation research, 100(6):782–794, 2007.

29. Mathieu Laplante and David M Sabatini. mtor signaling in growth control and disease. cell, 149(2):274–293, 2012.

30. Gaëlle Laurent, Florence Solari, Bogdan Mateescu, Melis Karaca, Julien Castel, Brigitte Bourachot, Christophe Magnan, Marc Billaud, and Fatima Mechta-Grigoriou. Oxidative stress contributes to aging by enhancing pancreatic angiogenesis and insulin signaling. Cell metabolism, 7(2):113–124, 2008.

31. LJ Lawson, VH Perry, and S Gordon. Turnover of resident microglia in the normal adult mouse brain. Neuroscience, 48(2):405–415, 1992.

32. Siu Sylvia Lee, Raymond YN Lee, Andrew G Fraser, Ravi S Kamath, Julie Ahringer, and Gary Ruvkun. A systematic rnai screen identifies a critical role for mitochondria in c. elegans longevity. Nature genetics, 33(1):40–48, 2003.

33. Arthur Liberzon, Aravind Subramanian, Reid Pinchback, Helga Thorvaldsdóttir, Pablo Tamayo, and Jill P Mesirov. Molecular signatures database (msigdb) 3.0. Bioinformatics, 27(12):1739–1740, 2011.

34. Carlos López-Otín, Maria A Blasco, Linda Partridge, Manuel Serrano, and Guido Kroemer. The hallmarks of aging. Cell, 153(6):1194–1217, 2013.

35. Lacy E Lowry and William A Zehring. Potentiation of natural killer cells for cancer immunotherapy: a review of literature. Frontiers in immunology, 8:1061, 2017.

36. Yasushi Magami, Takeshi Azuma, Hideto Inokuchi, Shinichiro Kokuno, Fuminori Moriyasu, Keiichi Kawai, and Takanori Hattori. Cell proliferation and renewal of normal hepatocytes and bile duct cells in adult mouse liver. Liver, 22(5):419–425, 2002.

37. Arulmani Manavalan, Manisha Mishra, Lin Feng, Siu Kwan Sze, Hiroyasu Akatsu, and Klaus Heese. Brain site-specific proteome changes in aging-related dementia. Experimental & molecular medicine, 45(9):e39–e39, 2013.

38. Barsanjit Mazumder, Prabha Sampath, Vasudevan Seshadri, Ratan K Maitra, Paul E DiCorleto, and Paul L Fox. Regulated release of l13a from the 60s ribosomal subunit as a mechanism of transcript-specific translational control. Cell, 115(2):187–198, 2003.

39. Ron Milo, Paul Jorgensen, Uri Moran, Griffin Weber, and Michael Springer. Bionumbers—the database of key numbers in molecular and cell biology. Nucleic acids research, 38(suppl_1):D750–D753, 2009.

40. Encarnacion Montecino-Rodriguez, Beata Berent-Maoz, and Kenneth Dorshkind. Causes, consequences, and reversal of immune system aging. The Journal of clinical investigation, 123(3):958–965, 2013.

41. Neda S Mousavy-Gharavy, Bryn Owen, Stephen J Millership, Pauline Chabosseau, Grazia Pizza, Aida Martinez-Sanchez, Emirhan Tasoez, Eleni Georgiadou, Ming Hu, Nicholas HF Fine, et al. Sexually dimorphic roles for the type 2 diabetes-associated c2cd4b gene in murine glucose homeostasis. bioRxiv, 2020.

42. Teresa Niccoli and Linda Partridge. Ageing as a risk factor for disease. Current biology, 22(17):R741–R752, 2012.

43. Janko Nikolich-Žugich. The twilight of immunity: emerging concepts in aging of the immune system. Nature immunology, 19(1):10–19, 2018.

44. Alessandro Ori, Brandon H Toyama, Michael S Harris, Thomas Bock, Murat Iskar, Peer Bork, Nicholas T Ingolia, Martin W Hetzer, and Martin Beck. Integrated transcriptome and proteome analyses reveal organ-specific proteome deterioration in old rats. Cell Systems, 1(3):224–237, 2015.

45. David Papadopoli, Karine Boulay, Lawrence Kazak, Michael Pollak, Frédérick Mallette, Ivan Topisirovic, and Laura Hulea. mtor as a central regulator of lifespan and aging. F1000Research, 8, 2019.

46. Marjolein J Peters, Roby Joehanes, Luke C Pilling, Claudia Schurmann, Karen N Conneely, Joseph Powell, Eva Reinmaa, George L Sutphin, Alexandra Zhernakova, Katharina Schramm, et al. The transcriptional landscape of age in human peripheral blood. Nature communications, 6:8570, 2015.

47. Daniel A Petkovich, Dmitriy I Podolskiy, Alexei V Lobanov, Sang-Goo Lee, Richard A Miller, and Vadim N Gladyshev. Using dna methylation profiling to evaluate biological age and longevity interventions. Cell metabolism, 25(4):954–960, 2017.

48. Simone Picelli, Omid R Faridani, Åsa K Björklund, Gösta Winberg, Sven Sagasser, and Rickard Sandberg. Full-length rna-seq from single cells using smart-seq2. Nature protocols, 9(1):171, 2014.

49. Angela Oliveira Pisco, Nicholas Schaum, Aaron McGeever, Jim Karkanias, Norma F Neff, Spyros Darmanis, Tony Wyss-Coray, Stephen R Quake, et al. A single cell transcriptomic atlas characterizes aging tissues in the mouse. bioRxiv, page 661728, 2019.

50. Noa Rappaport, Noam Nativ, Gil Stelzer, Michal Twik, Yaron Guan-Golan, Tsippi Iny Stein, Iris Bahir, Frida Belinky, C Paul Morrey, Marilyn Safran, et al. Malacards: an integrated compendium for diseases and their annotation. Database, 2013, 2013.

51. Noa Rappaport, Michal Twik, Noam Nativ, Gil Stelzer, Iris Bahir, Tsippi Iny Stein, Marilyn Safran, and Doron Lancet. Malacards: A comprehensive automatically-mined database of human diseases. Current Protocols in Bioinformatics, 47(1):1–24, 2014.

52. Noa Rappaport, Michal Twik, Inbar Plaschkes, Ron Nudel, Tsippi Iny Stein, Jacob Levitt, Moran Gershoni, C Paul Morrey, Marilyn Safran, and Doron Lancet. Malacards: an amalgamated human disease compendium with diverse clinical and genetic annotation and structured search. Nucleic acids research, 45(D1):D877–D887, 2017.

53. Uku Raudvere, Liis Kolberg, Ivan Kuzmin, Tambet Arak, Priit Adler, Hedi Peterson, and Jaak Vilo. g: Profiler: a web server for functional enrichment analysis and conversions of gene lists (2019 update). Nucleic acids research, 47(W1):W191–W198, 2019.

54. Richard D Riley, Julian PT Higgins, and Jonathan J Deeks. Interpretation of random effects meta-analyses. Bmj, 342, 2011.

55. Nicholas Schaum, Benoit Lehallier, Oliver Hahn, Róbert Pálovics, Shayan Hosseinzadeh, Song E Lee, Rene Sit, Davis P Lee, Patricia Morán Losada, Macy E Zardeneta, et al. Ageing hallmarks exhibit organ-specific temporal signatures. Nature, pages 1–7, 2020.

56. Roland G Scollay, Eugene C Butcher, and Irving L Weissman. Thymus cell migration: quantitative aspects of cellular traffic from the thymus to the periphery in mice. European journal of immunology, 10(3):210–218, 1980.

57. Giovanni Stallone, Barbara Infante, Concetta Prisciandaro, and Giuseppe Grandaliano. mtor and aging: An old fashioned dress. International journal of molecular sciences, 20(11):2774, 2019.

58. Fiona A Stewart, Juliana Denekamp, and DG Hirst. Proliferation kinetics of the mouse bladder after irradiation. Cell Proliferation, 13(1):75–89, 1980.

59. Aravind Subramanian, Pablo Tamayo, Vamsi K Mootha, Sayan Mukherjee, Benjamin L Ebert, Michael A Gillette, Amanda Paulovich, Scott L Pomeroy, Todd R Golub, Eric S Lander, et al. Gene set enrichment analysis: a knowledge-based approach for interpreting genome-wide expression profiles. Proceedings of the National Academy of Sciences, 102(43):15545–15550, 2005.

60. Robi Tacutu, Daniel Thornton, Emily Johnson, Arie Budovsky, Diogo Barardo, Thomas Craig, Eugene Diana, Gilad Lehmann, Dmitri Toren, Jingwei Wang, et al. Human ageing genomic resources: new and updated databases. Nucleic acids research, 46(D1):D1083–D1090, 2017.

61. Monica Teta, Simon Y Long, Lynn M Wartschow, Matthew M Rankin, and Jake A Kushner. Very slow turnover of *β*-cells in aged adult mice. Diabetes, 54(9):2557–2567, 2005.

62. John Tower. Programmed cell death in aging. Ageing research reviews, 23:90–100, 2015.

63. Jan Vijg and Yousin Suh. Genome instability and aging. Annual review of physiology, 75:645–668, 2013.

64. Thomas Weichhart. mtor as regulator of lifespan, aging, and cellular senescence: a mini-review. Gerontology, 64(2):127–134, 2018.

65. Liset Westera, Julia Drylewicz, Ineke Den Braber, Tendai Mugwagwa, Iris Van Der Maas, Lydia Kwast, Thomas Volman, Elise HR Van De Weg-Schrijver, István Bartha, Gerrit Spierenburg, et al. Closing the gap between t-cell life span estimates from stable isotope-labeling studies in mice and humans. Blood, The Journal of the American Society of Hematology, 122(13):2205–2212, 2013.

66. Methodios Ximerakis, Scott L Lipnick, Sean K Simmons, Xian Adiconis, Brendan T Innes, Danielle Dionne, Lan Nguyen, Brittany A Mayweather, Ceren Ozek, Zachary Niziolek, et al. Single-cell transcriptomics of the aged mouse brain reveals convergent, divergent and unique aging signatures. bioRxiv, page 440032, 2018.

67. Xiang Zhou, Wen-Juan Liao, Jun-Ming Liao, Peng Liao, and Hua Lu. Ribosomal proteins: functions beyond the ribosome. Journal of molecular cell biology, 7(2):92–104, 2015.

## References

1. Jacob C Kimmel, Lolita Penland, Nimrod D Rubinstein, David G Hendrickson, David R Kelley, and Adam Z Rosenthal. Murine single-cell rna-seq reveals cell-identity-and tissue-specific trajectories of aging. Genome research, 29(12):2088–2103, 2019.

2. Monika S Kowalczyk, Itay Tirosh, Dirk Heckl, Tata Nageswara Rao, Atray Dixit, Brian J Haas, Rebekka K Schneider, Amy J Wagers, Benjamin L Ebert, and Aviv Regev. Single-cell rna-seq reveals changes in cell cycle and differentiation programs upon aging of hematopoietic stem cells. Genome research, 25(12):1860–1872, 2015.

3. Andreas Krämer, Jeff Green, Jack Pollard Jr, and Stuart Tugendreich. Causal analysis approaches in ingenuity pathway analysis. Bioinformatics, 30(4):523–530, 2013.

4. Nicholas Schaum, Benoit Lehallier, Oliver Hahn, Róbert Pálovics, Shayan Hosseinzadeh, Song E Lee, Rene Sit, Davis P Lee, Patricia Morán Losada, Macy E Zardeneta, et al. Ageing hallmarks exhibit organ-specific temporal signatures. Nature, pages 1–7, 2020.

5. Robi Tacutu, Daniel Thornton, Emily Johnson, Arie Budovsky, Diogo Barardo, Thomas Craig, Eugene Diana, Gilad Lehmann, Dmitri Toren, Jingwei Wang, et al. Human ageing genomic resources: new and updated databases. Nucleic acids research, 46(D1):D1083–D1090, 2017.

